# Transcriptome landscape of a thermal-tolerant coral endosymbiont reveals molecular signatures of symbiosis and dysbiosis

**DOI:** 10.1101/508184

**Authors:** Anthony J. Bellantuono, Katherine E. Dougan, Camila Granados-Cifuentes, Mauricio Rodriguez-Lanetty

## Abstract

**Background:** Reef-building corals depend upon a nutritional endosymbiosis with photosynthetic dinoflagellates (family Symbiodiniaceae) for the vast majority of their energetic needs. While this mutualistic relationship is impacted by numerous stressors, heat stress from warming oceans is a predominant threat to coral reefs, placing the future of the world’s reefs in peril due to climate change. While some Symbiodiniaceae species exhibit tolerance to thermal stress, the molecular basis for this tolerance is largely unexplored. To identify the underpinnings of heat tolerance and symbiosis, we compared the *in hospite* and free-living transcriptomes of *Durusdinium trenchii*, a pan-tropical heat-tolerant Symbiodiniaceae, under stable temperature conditions and acute hyperthermal stress.

**Results:** We discovered that under stable conditions, *in hospite* cells exhibited lower transcriptional activity than free-living counterparts, suggesting the shutdown of genes uniquely required for a free-living lifestyle in a process we term transcriptome reduction. However, under hyperthermal stress the transcriptional response was larger *in hospite* than in the free-living state, indicating an exacerbated stress environment within the host cell. While we identified a core heat stress response shared between free-living and *in hospite D. trenchii*, this represents only a minority of the stress-induced transcriptional activity. Our findings also revealed distinct processes at work in free-living and *in hospite* cells under non-stress conditions, with free-living *D. trenchii* exhibiting evidence of sexual reproduction. We also found that *in hospite* cells display putative host evasion activity including components of apicomplexan parasite invasion machinery.

**Conclusions:** In this work, we unraveled the molecular signatures of symbiont heat tolerance within the host, which is a critical step to enable the development of engineered endosymbionts as a tool for restoration of coral reefs. Our findings also highlight the necessity of conducting experiments on Symbiodiniaceae in the appropriate symbiotic state, as the stress responses of free-living cultures may not be representative of Symbiodiniaceae within a holobiont. We suggest that the necessity of an intermittent free-living state, outside of its highly co-evolved symbiosis, prevents genome reduction as seen in obligate endosymbionts. We posit that transcriptomic reduction of non-essential genes *in hospite* may be an alternative adaptive strategy for endosymbionts in ancient symbiotic assemblages.

## Introduction

Reef-building corals engage in an obligate endosymbiosis with a diverse group of dinoflagellate algae in the family Symbiodiniaceae, which is composed of seven recently described genera (LaJeunesse et al., 2018). In this nutritional symbiosis, the algal partner transfers up to 90% of the carbon it fixes to the coral host (Muscatine et al., 1984). However, this critical mutualism is sensitive to environmental stressors and can break down in a process termed coral bleaching (Glynn, 1993), most commonly due to elevated seawater temperature linked to anthropogenic climate change (Hoegh-Guldberg, 1999). High temperature anomalies are continuing to intensify, with 2015–2016 exhibiting the most intense global bleaching event on record (Eakin et al., 2016; Hughes et al., 2017). Recent models forecast that mass bleaching will be an annual event before 2050 (Maynard et al., 2017). Even if dramatic international actions manage to limit warming to 1.5 °C, coral populations are predicted to remain in peril (Anthony et al., 2017).

There is an existing body of work demonstrating that the shift of the dominant symbiont (shuffling) or acquisition of new species of symbionts (switching) from the environment enables holobionts, the coral host and its diverse assemblage of symbionts, to better withstand climate-related increases in seawater temperatures (Baker, 2001; Berkelmans and Van Oppen, 2006; Jones et al., 2008; Rowan, 2004; Stat et al., 2006). More recently, significant differences in heat tolerance have been reported both within a single Symbiodiniaceae genus (Díaz-Almeyda et al., 2017) as well as between genera (Grégoire et al., 2017), revealing that different Symbiodiniaceae species have differing thermal optima and tolerances. A well-studied case is *Durusdinium trenchii* (previously *Symbiodinium trenchii* [LaJeunesse et al., 2018]), an ITS2 – D1a subtype within Clade D that has been shown to confer increases in heat tolerance of between 1-1.5 °C to coral hosts (Berkelmans and Van Oppen, 2006; Rowan, 2004). This heat tolerance is also observed in culture conditions; *D. trenchii* can maintain maximum photosynthetic rates at elevated temperatures in comparison to the majority of other investigated Symbiodiniaceae species (Grégoire et al., 2017). However, alternate heat-resistant assemblages of host and symbiont come with a cost, showing decreases in growth rates (Jones and Berkelmans, 2011; Little et al., 2004).

The potential benefits to corals hosting *D. trenchii* in response to increasingly warm ocean conditions is not only evident in Pacific reef corals (Baker, 2003), where this species of symbiont is genetically diverse and is thought to have originated, but also in the Caribbean Sea, where it is thought to have invaded in the past half century (Pettay et al., 2015). Silverstein et al. (2017) have demonstrated that the Caribbean scleractinian coral *Montastraea cavernosa* colonized by *D. trenchii* persist *in hospite* on thermally stressed reefs, whereas conspecific corals hosting type C3 symbionts underwent massive bleaching.

Due to the current ecological crisis facing corals, there has been a growing movement to develop purposeful interventions to increase the thermal tolerance of many heat-sensitive coral holobionts, including experimental evolution and interspecific hybridizations (Chakravarti et al., 2017; Chan et al., 2018). Recently, the use of genome editing approaches has been proposed, with the primary goal of engineering heat-sensitive endosymbionts in order to produce more heat-tolerant coral holobionts (Levin et al., 2017). This requires the characterization of molecular mechanisms used by heat-resistant Symbiodiniaceae species so that taxa exhibiting less resistant phenotypes can be genetically engineered into heat-resistant symbionts.

By using a transcriptomic approach, we aimed to characterize the mRNA profiles associated with the response to thermal stress in the heat-resistant *D. trenchii*. We assessed the gene expression response of the symbiont both in the free-living state and *in hospite* when stably associated with a cnidarian host. This design allowed us to gauge the effect of symbiosis on the molecular response of the symbiont to heat stress. In order to achieve sufficient sequencing depth of *in hospite D. trenchii*, we implemented a novel symbiont mRNA enrichment technique. This powerful comparative approach also facilitated an examination of the molecular changes underlying the transition from a free-living to symbiotic state. This represents the first work to characterize the transcriptional modifications associated with *in hospite* and free-living states of a Symbiodiniaceae, and the first to characterize the divergent stress responses between the symbiotic and free-living environment in a very ecologically important coral symbiont, *D. trenchii*.

Our investigation reveals the dramatic differences in gene expression between symbiotic and free-living *D. trenchii*. Further, we show that these differences are magnified under conditions of thermal stress. When D. trenchii are in symbiosis, we note a significant downregulation of transcripts. This molecular signature suggests that the host provides a stabilized environment for *D. trenchii* in contrast to the free-living state. Under thermal stress, the more active transcriptomic response of the dinoflagellate *in hospite* in comparison to the free-living state suggests that the dinoflagellates are responding to a less hospitable environment. This suggests that the dinoflagellates living within host cells are responding to a more inhospitable environment under thermal stress as disturbed host cells may intensify the stressful state of the symbiont. This result brings critical attention to the necessity of examining thermal stress responses of coral endosymbionts *in hospite* in order to capture the functional profile actually relevant to coral holobiont persistence. Studies based solely on free-living cultures provide a biased perspective that is confounded by responses to an external environment markedly different from the holobiont.

## Results and Discussion

### Transcriptome Analysis

A reference transcriptome of 82,273 transcripts was assembled, filtered for non-Symbiodiniaceae transcripts, and used as the basis for differential expression analysis (NCBI TSA BioProject PRJNA508937). Principal component analysis (PCA) of all analyzed transcripts shows dramatic separation of treatments by both symbiotic state as well as thermal stress (Fig. 2*A*). PC1, the principal component separating *in hospite* and free-living treatments, is driven by 11,933 transcripts and explains 40.5% of the variance, whereas PC2, the axis separating thermal treatment, is driven by 11,904 transcripts and explains 15.5% of the variance.

**Figure 1.**
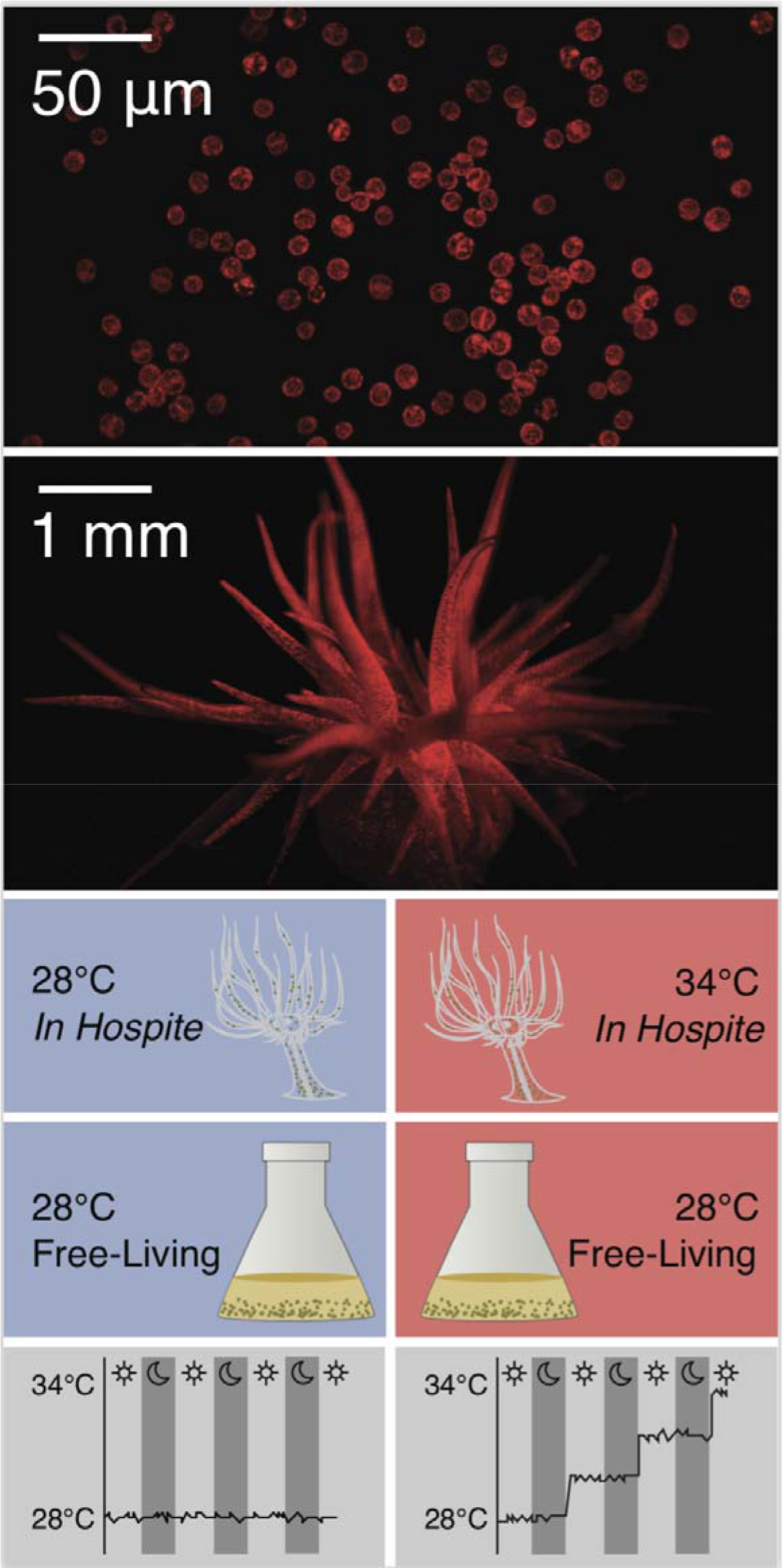
Durusdinium trenchii in free-living culture state (top) and in hospite in E. pallida (second panel). Schematic of experimental treatments (lower panels) comparing symbiotic state and thermal stress, with temperature and light profiles.

**Figure 2.**
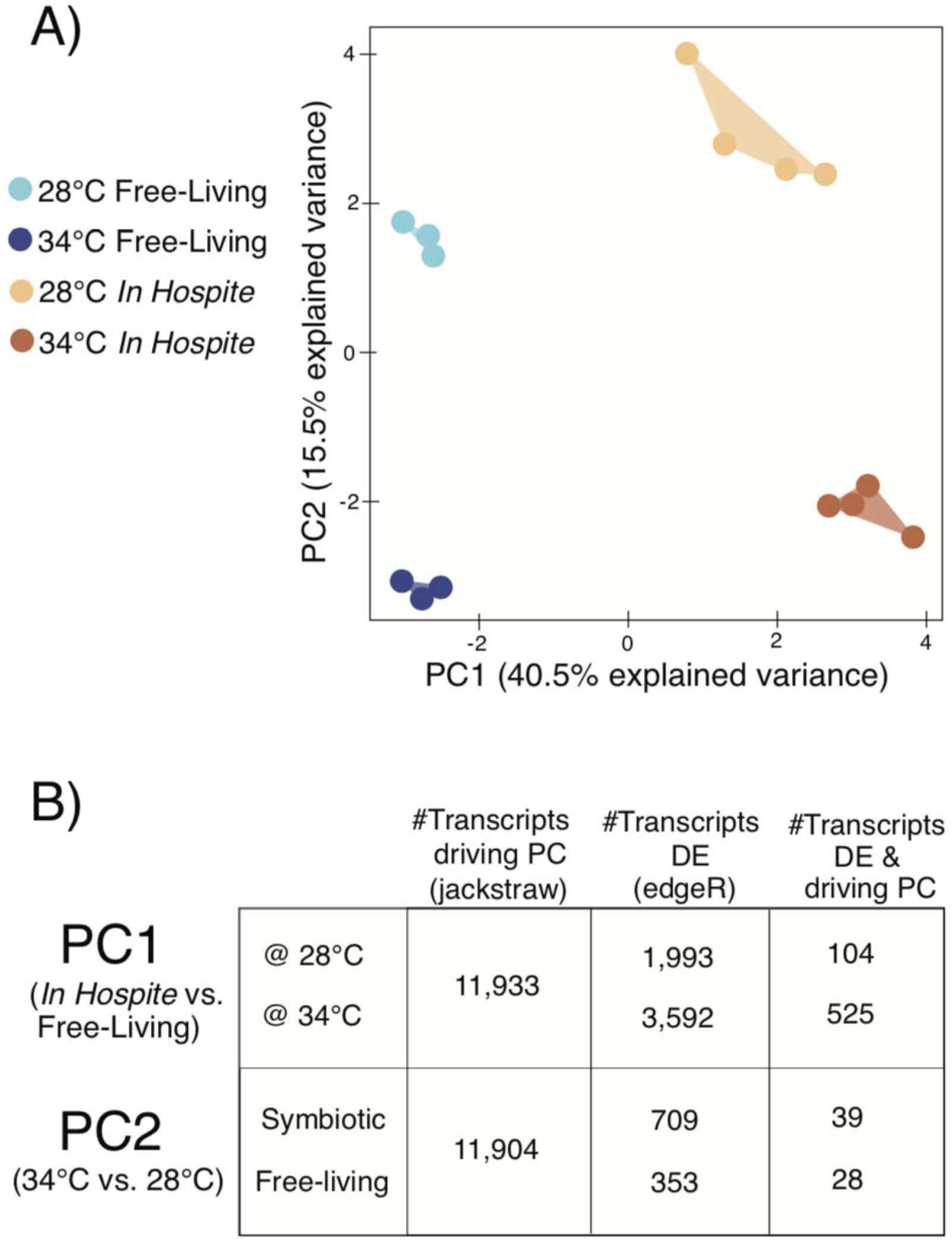
A) Principal component analysis depicting the ordination of biological replicates by all analyzed transcripts. (B) Summary of the number of transcripts driving PC1 and PC2, the number of transcripts DE according to edgeR, and the number of transcripts that were both DE and significant drivers of PC1 and PC2.

Statistical analysis for the identification of differentially expressed transcripts (FDR corrected p-value < 0.01, greater than log_2_ 2-fold difference) reveals that the *in hospite* condition in *D. trenchii* is marked by a substantial downregulation of gene expression, with 368 transcripts upregulated versus 1625 downregulated at the 28°C (Fig. 3A). This larger level of downregulation *in hospite* is maintained under the heat stress comparison (34°C), with 2799 transcripts downregulated versus 793 upregulated in comparison to the free-living state (Fig. 3A). Notably, the *in hospite* response to heat stress is much more pronounced than that in the free-living condition, with a total of 709 transcripts differentially regulated as a function of thermal stress *in hospite* versus 353 transcripts in the free-living condition (Fig. 3B). Out of the 709 transcripts involved in the *in hospite* response to heat, 624 transcripts (365 upregulated and 259 downregulated) were specific to the *in hospite* condition as these transcripts were not differentially expressed in response to thermal challenge in free-living *D. trenchii* (Fig. 3B). The transcriptional response to thermal stress shared between free-living and *in hospite D. trenchii* represented only 12% of the heat-modulated transcriptional response *in hospite*. This shared response could be considered the core heat response, with activity regardless of the symbiotic state of the algal endosymbiont. However, this represents a minority of detected transcriptional activity. The majority of transcripts involved in response to heat stress depend upon the symbiotic condition of the dinoflagellate (e.g. whether the cell is *in hospite* or free-living). Our findings demonstrate that *D. trenchii* has distinct transcriptomic profiles *in hospite* and in free-living states, and further, that the transcriptomic heat stress response is vastly different between *in hospite* and free-living conditions. This brings critical attention to the importance of examining the response of Symbiodiniaceae holistically, as the responses in culture may not be representative of *in hospite* endosymbionts. To date, the vast majority of transcriptomic assessments of Symbiodiniaceae within the context of thermal stress have been performed exclusively on free-living cultures (Barshis et al., 2014; Baumgarten et al., 2013; Díaz-Almeyda et al., 2017; Gierz et al., 2017; Levin et al., 2016; Rachel Ashley Levin et al., 2017; Liew et al., 2017; Xiang et al., 2015), and while they have been insightful, they may present a reduced transcriptional picture of the alterations that occur in Symbiodiniaceae *in hospite*. Only one other investigation directly compared gene expression *in hospite* to free-living conditions (Rosic et al., 2010), but it focused on only two heat-shock protein genes. We propose that it is imperative to consider the *in hospite* environment in the assessment of Symbiodiniaceae response to thermal stress, and that culture-based studies require careful consideration when extending findings to *in hospite* conditions.

**Figure 3.**
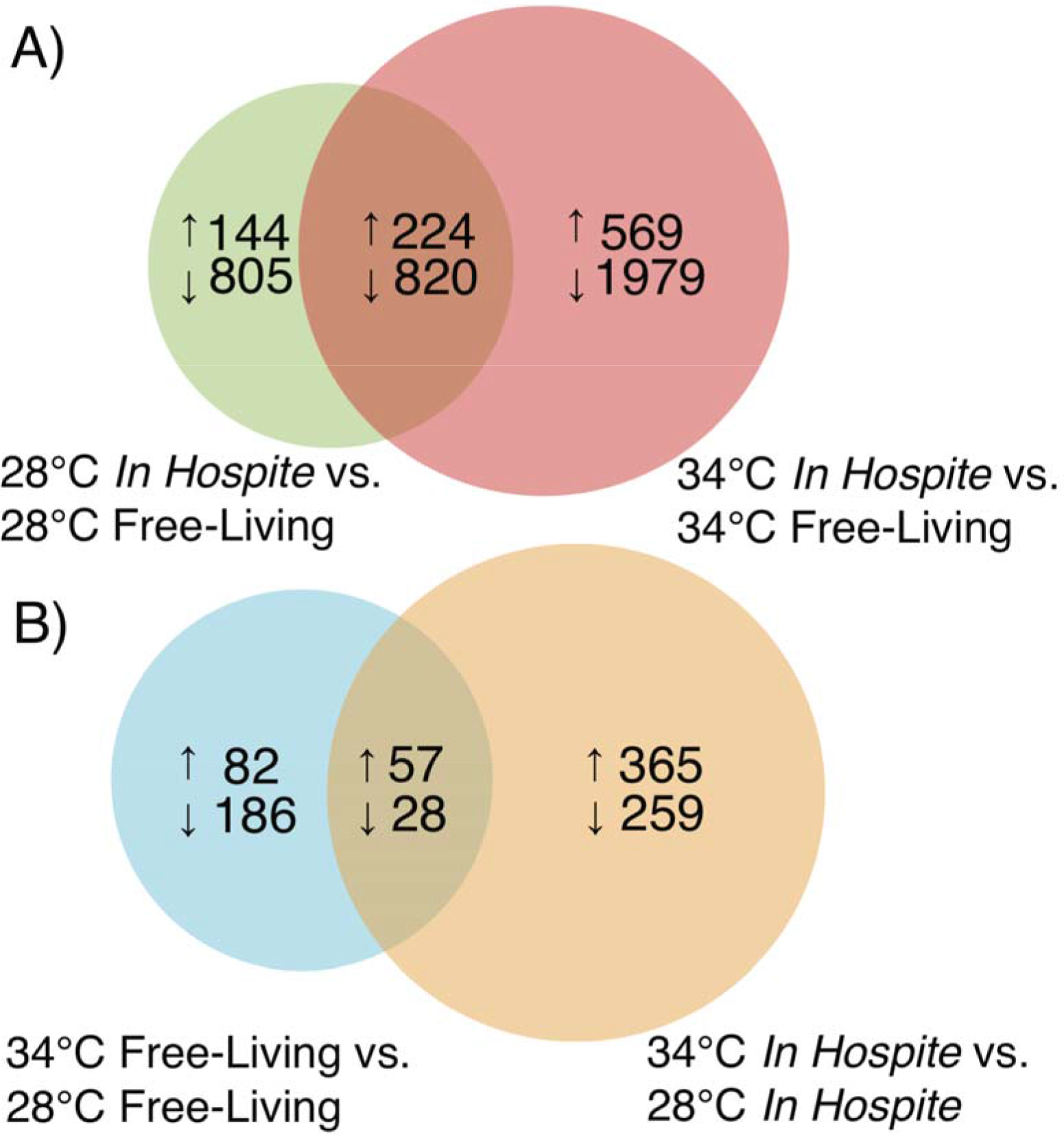
Venn diagrams of the number of differentially expressed transcripts by pairwise comparison (FDR-corrected p < 0.01, greater than 2-fold difference). A) Contrasts transcriptional changes between in hospite and free-living transcripts at 28°C and 34°C. B) Comparison of transcripts differentially expressed in the free-living and in hospite condition, with the core heat stress response represented by the overlapping region.

### A Small Core Thermal Stress Response is Shared Between Free-living and *in hospite D. trenchii*

Within the conserved transcriptional changes under thermal stress shared between the free-living and *in hospite* states of *D. trenchii* (Fig. 3, Fig. 4), 57 transcripts were upregulated and 28 downregulated. Examination at the transcript level reveals that the magnitude of expression within these core transcripts is, overall, higher in free-living *D. trenchii* than *in hospite* (Fig. 4). With a relatively small number of transcripts representing the core response, we were not able to perform meaningful GO term enrichment analysis. Instead, our interpretation of this small conserved core response is based upon transcript annotations.

**Figure 4.**
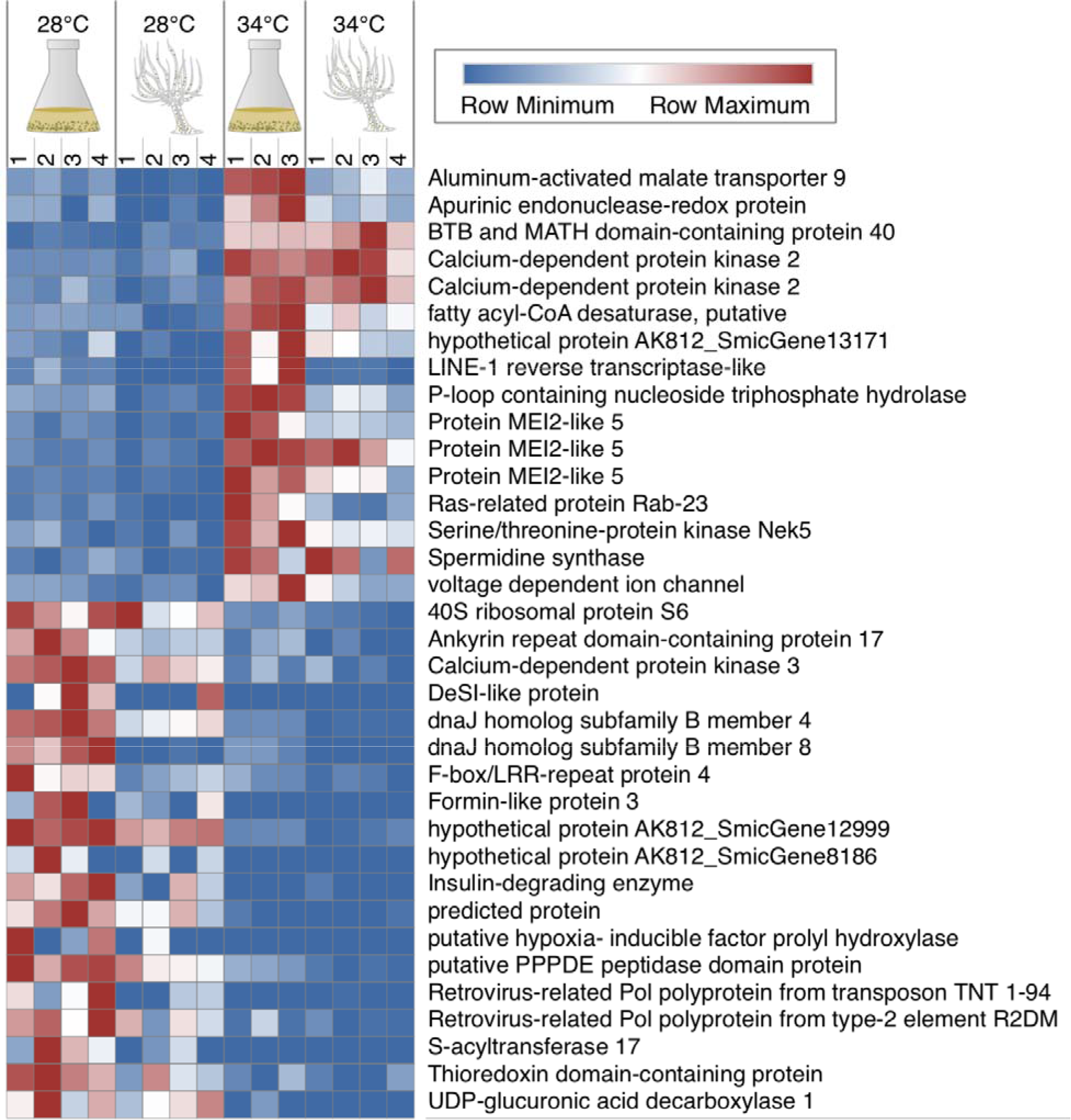
Core heat stress response transcripts. The heatmap reflects annotated transcripts differentially expressed in both in hospite and free-living thermal stress conditions. (Unannotated transcripts omitted.)

Though the core set of transcripts differentially expressed in response to thermal stress (Fig. 3) represents the minority of transcriptional alterations under stress *in hospite* and in free-living cells, it sheds light on key processes conserved in the *D. trenchii* stress response independent of symbiotic state. These processes include modifications to membrane fluidity, DNA repair, anion channel activity, spermidine synthase upregulation, and transposition. With their indispensability in both the *in hospite* and free-living stress responses, these associated transcripts are potentially important candidates in the identification of the genetic basis for thermal tolerance in *D. trenchii*.

Maintenance of membrane viscosity is critical for cell homeostasis as hyperthermal stress increases the fluidity of membranes, potentially leading to disintegration of the lipid bilayer (Allakhverdiev et al., 2008). In Symbiodiniaceae, thylakoid membrane lipid composition is a key determinant of thermal-stress sensitivity (Tchernov et al., 2004). Within the core heat stress response, we identify an ortholog to a *fatty acyl-CoA desaturase*, an enzyme involved in the remodeling of cell membranes for homeoviscous adaptation (Hazel, 1995) that is potentially critical in modifying fluidity to accommodate increasing temperatures.

The upregulation of *apurinic endonuclease-redox protein* (*ARP*) both *in hospite* and in free-living conditions suggests that repair to oxidative DNA base damage occur in both conditions. The apurinic/apyrimidinic (AP) endonucleases are important DNA repair enzymes involved in two overlapping pathways: DNA glycosylase-initiated base excision and AP endonuclease-initiated nucleotide incision repair (Akishev et al., 2016). In the context of heat stress, this may allow *D. trenchii* a faster response to DNA damage than other Symbiodiniaceae species.

Anion channels, spermidine synthase, and retrotransposons are disparate entities which can each serve as heat stress response mediators with many downstream effectors. In plants, anion channels are a master regulator of stress response (reviewed by Rob et al., 2012). In the core heat stress response, we find the upregulation of a transcript putatively coding for aluminum-activated malate transporter 9 (ALMT9). In plants, ALMT9 is involved in the release of chloride currents in guard cells and malate homoeostasis. In response to stress, the release of chloride and malate leads to the rapid stomatal closure (Dreyer et al., 2012). Though the precise role of ALMT9 in stress response in Symbiodiniaceae species is unknown, it may be involved in downstream stress response signaling. A transcript coding for spermidine synthase is also upregulated across heat stress treatments. Spermidine synthase in *Arabidopsis* dramatically improves stress tolerance when overexpressed via its effect on multiple downstream targets (Kasukabe et al., 2004), while exogenous application of the protein increases antioxidant tolerance in creeping bentgrass (Li et al., 2015). An additional transcript of the core response potentially acting as an upstream effector of numerous targets codes for LINE-1 reverse transcriptase. Recent work by Chen et al. (2018) identified Ty1-copia-type LTR retrotransposons in *Symbiodinium microadriaticum* which are activated by heat stress, suggesting that this may be a mechanism for the generation of genetic variation under heat stress. Diatoms also possess stress-activated retrotransposons, which are postulated to serves as generators of genetic variability (Maumus et al., 2009). The involvement of retrotransposons in stress response has also been observed across taxa. In plants, retrotransposons are activated by stress and go on to activate numerous host regulatory pathways (Grandbastien, 2015, 1998).

### *D. trenchii* response to heat stress depends on the symbiotic state

The majority (88% *in hospite* and 76% free-living) of the transcriptional response to thermal stress is specific to the symbiotic state of the dinoflagellate (Fig. 3B). Both free-living and *in hospite D. trenchii* have distinct responses to a thermal challenge of 34°C not only at the level of transcript identity (Fig. 3), but also with regard to enriched gene ontology terms which are unique to each symbiotic state (Fig. 5). *In hospite D. trenchii* have transcriptional activity with parent GO terms such as mRNA transport, N-terminal protein amino acid acetylation, and cellular response to temperature stimulus, all of which are not enriched in the free-living transcriptomic response to heat stress (Fig. 5). There is also a striking *in hospite-*specific decrease in GO terms within the category “binding of sperm to zona pellucida” under heat stress. In the case of the free-living state, negative regulation of viral genome replication is highly downregulated in contrast to the symbiotic state (Fig. 5). While the analysis reveals that the majority of ontologies affected by heat stress are disparate depending upon symbiotic state, decreased activity of vesicle-mediated transport is a common theme in the heat stress response of both *in hospite* and free-living *D. trenchii* (Fig. 5). Despite this particular instance of overlap in function indicated by gene ontology analysis, the transcripts underlying vesicle-mediated transport in free-living and symbiotic treatments are distinct in the symbiotic and free-living conditions (Supplement_2_DEGs_Annotations.xlsx).

**Figure 5.**
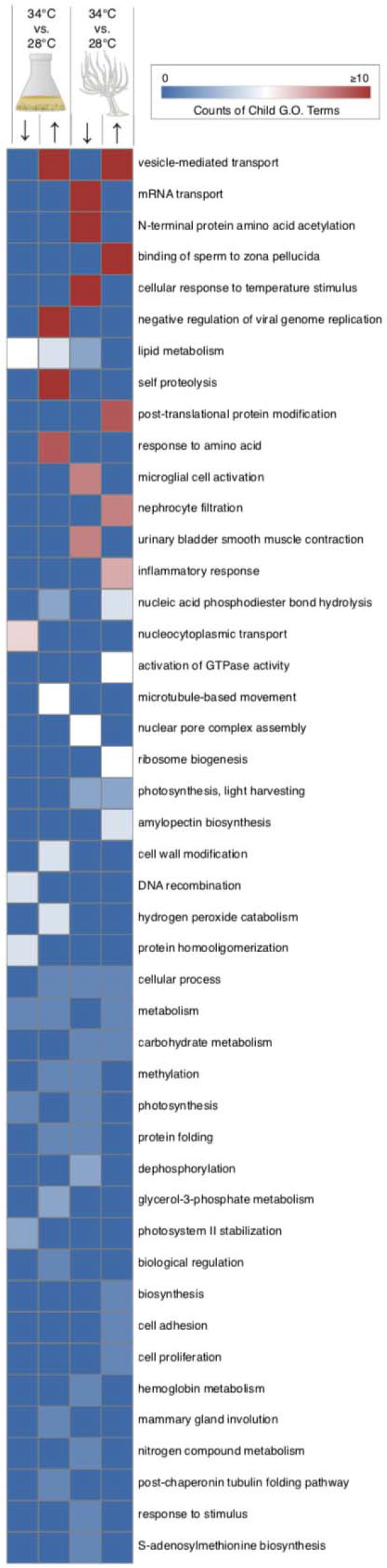
Counts of GO terms enriched exclusively in the free-living and in hospite heat stress response. (Terms associated with transcripts differentially expressed in both in hospite and free-living conditions were excluded from this enrichment analysis).

Remarkably, despite the more highly responsive transcriptome of *in hospite D. trenchii* to heat stress than free-living cells, free-living *D. trenchii* mount a more dramatic response to photoxidative stress than *in hospite* at both 28°C and 34°C (Fig. 6). Gene ontology terms for thermoception and response to oxidative stress are also highly represented in free-living *D. trenchii* at 34°C, whereas no enrichment is detected *in hospite* (Fig. 6). This suggests that the intracellular environment of the host provides some level of protection for the endosymbiont from some aspects of photothermal stress, potentially by a modified light environment within the host. This is supported by earlier work by Bhagooli and collaborators (Bhagooli et al., 2008), who found that multiple genera of Symbiodiniaceae exposed to photothermal stress *in hospite* had unaffected photosynthetic output and less impact to maximum quantum yield in comparison to cultured cells across genera. Coral hosts possess multiple mechanisms for sheltering endosymbionts including photoprotective host chromoproteins (Dove et al., 2001; Quick et al., 2018; Salih et al., n.d.; Smith et al., 2013) and other host-mediated modifications to the internal light environment, such as scattering by the coral skeleton (Enríquez et al., 2005; Wangpraseurt et al., 2012). While this sheltering *in hospite* may, on its surface, appear contradictory, we posit that the *in hospite* state has a more complex transcriptional response to thermal stress overall, but that the host may reduce light stress.

**Figure 6.**
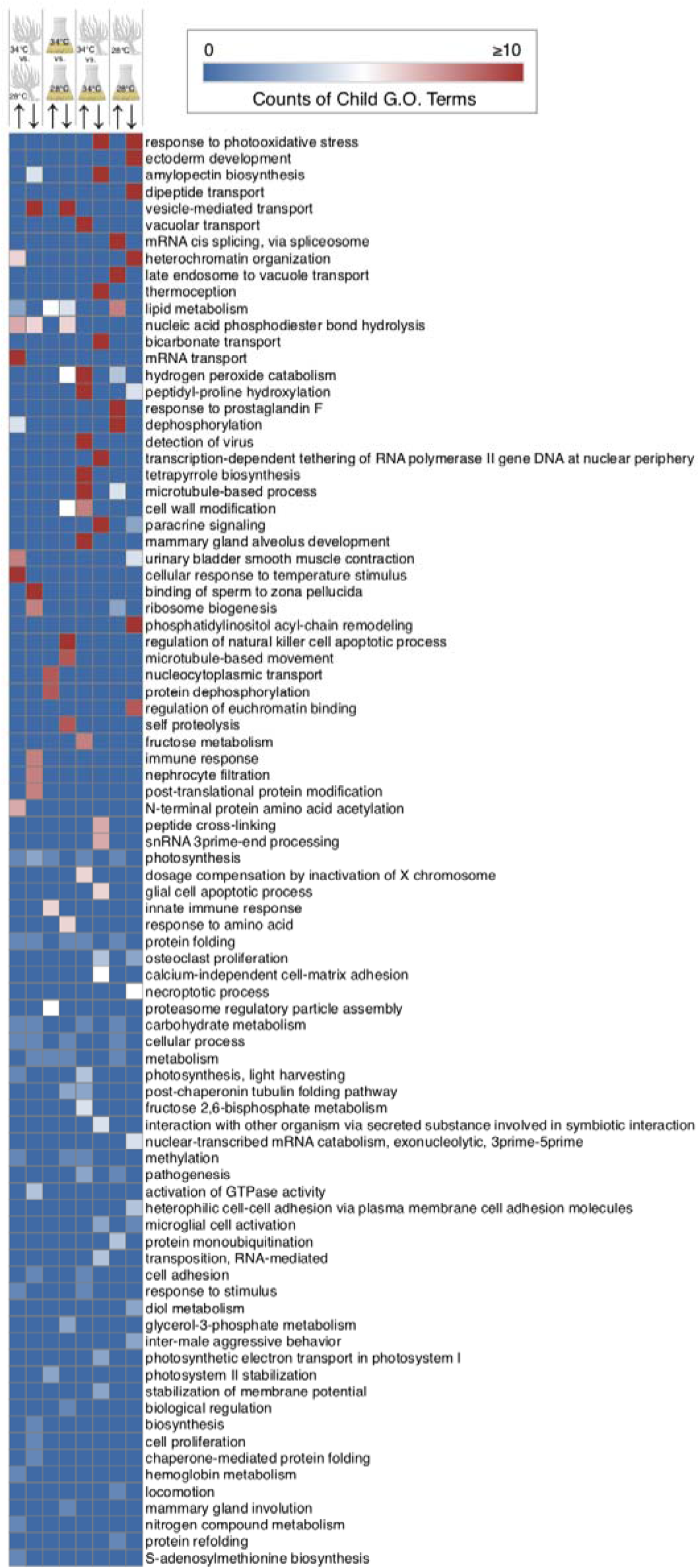
Counts of GO terms associated with up- and downregulated transcripts across all comparisons. Parent GO terms are listed for all categories with a minimum sum of two daughter terms across all comparisons.

By examining ontologies from differentially expressed transcripts exclusively associated with free-living and *in hospite* treatments, we found that the *in hospite* heat stress response is highly enriched for mRNA transport and cellular response to temperature stimulus (Fig. 5). We suggest that host sheltering of *in hospite* cells light may contribute to a symbiont’s capacity to maintain some cellular functions which may be heavily impacted in unsheltered free-living cells. This may result in the mounting of a more effective transcriptional response to heat stress. Under acute thermal stress, yeast block mRNA transport of transcripts uninvolved in the heat stress response (Saavedra et al., 1996). If unsheltered free-living cells are under more severe stress, experiencing the unfiltered brunt of light in combination with heat stress, mRNA transport may be highly inhibited, as suggested by our analyses.

The *in hospite* state also exhibits a dramatic reduction in GO term enrichment within the biological process binding of sperm to zona pellucida, a response not observed in the free-living comparison (Fig. 5). One of the transcripts underlying this enrichment (TRINITY_DN30024_c0_g1_i6) has high identity to *sperm surface protein Sp17*, a protein possessed by sperm which binds the zona pellucida during mammalian fertilization, critical for cell adhesion (O’rand et al., 1995; Richardson et al., 1994). The expression of the putative *Sp17* ortholog may function in the mediation of normal symbiosis, potentially involved in adhesion between the symbiont and host, while the observed decrease in expression may be a part of dysbiosis.

Taken together, the intracellular environment experienced by *in hospite* symbionts represents a complex set of interactions between the host, the external environment, and the endosymbiont. The disparate transcriptomes of stress *in hospite* and free-living likely result from a combination of protection from some aspects of stress by the host, the complexity of an intracellular environment compared with that of a free-living cell, and further, the transcriptional activity involved in the mediation of the symbiosis, as discussed in sections that follow.

### Alterations required for the transition between free-living and symbiotic states

Comparing the *in hospite* and free-living transcriptomes of *D. trenchii* provides a window into the alterations necessary for symbiotic living. Several processes are exclusively enriched *in hospite* under non-stress conditions at 28°C, in comparison to the free-living state (28°C symbiotic vs. 28°C free-living) (Fig. 6). Late endosome to vacuole transport and mRNA cis splicing via spliceosome are both highly enriched *in hospite* along with response to prostaglandin F. The roles of increased lysosome activity and enhanced cis splicing in symbiosis are unknown, but complexities in Symbiodiniaceae mRNA splicing, including trans-splicing (Zhang et al., 2007) and non-canonical intron splicing (Aranda et al., 2016; Lin, 2011; Mendez et al., 2015). The enrichment of response to prostaglandin F is extremely curious as the highest concentrations of prostaglandins are found in corals, where these compounds are produced by the host (Valmsen et al., 2001; Weinheimer and Spraggins, 1969). Hydrogen peroxide catabolism is enriched *in hospite* at both 28°C and 34°C in comparison to the free-living state, with higher activity at 34°C. Increased scavenging of hydrogen peroxide *in hospite* could potentially be adaptive and necessary for maintaining stable symbiosis, as while the release of peroxide in the free-living state is likely of minimal detriment, the release of this form of ROS *in hospite* is held to induce an immune response on the part of the host (reviewed by Weis, 2008).

In addition to the aforementioned enrichment of photooxidative stress terms in the free-living state under thermal challenge (34°C), the transcripts upregulated in the free-living state at 28°C are also enriched for numerous terms associated with response to photooxidative stress. While this is amongst the highest activity of parent GO terms in the free-living condition at both thermal treatments, no increase in enrichment is found in the *in hospite* condition at either 28°C or 34°C. Dipeptide transport as well as heterochromatin organization are also highly enriched in the free-living condition at 28°C.

### *D. trenchii* exhibits transcriptome reduction *in hospite*

Genome reduction is a common phenomenon in obligate microbial endosymbionts (Martínez-Cano et al., 2015; McCutcheon and Moran, 2012) which can occur via multiple mechanisms. In obligate intracellular endosymbioses, there are some common demographic conditions imposed upon the endosymbiont, namely relatively small effective population size of endosymbionts, restriction to specific hosts, and limited recombination between endosymbiont strains within the host (Martínez-Cano et al., 2015). These conditions set the stage for genetic drift and loss of genes that are not essential *in hospite*. However, this appears to not be the case in coral endosymbiosis, given that all sequenced Symbiodiniaceae genomes are of approximately one gigabase in size (Aranda et al., 2016; Lin et al., 2015; Liu et al., 2018; Shoguchi et al., 2013), nor should it be expected, given that the coral endosymbiont must maintain the full complement of genes for free-living persistence, as the majority of corals uptake their endosymbionts from the free-living environment (Baird et al., 2009; Richmond and Hunter, 1990). Further, Symbiodiniaceae have been found to exist in a free-living state in numerous environments, including sediments and other benthic substrates, as epiphytes, and in the water column (Granados-Cifuentes et al., 2015; Littman et al., 2008; Takabayashi et al., 2012).

Despite this lack of genome reduction in Symbiodiniaceae, we identified a significant downregulation of gene expression *in hospite* compared to free-living *D. trenchii* at both temperatures. Free-living Symbiodiniaceae appear to require to a larger transcriptional repertoire to persist in an extracellular environment, for functions presumably including motility and coping with fluctuations experienced outside of the confines of a host cell. This substantial downregulation of transcripts *in hospite* is associated with enrichment of transcripts associated with heterochromatin organization histone H3-K9 methylation (Fig. S2). The hypermethylated state of cultured, free-living Symbiodinaceae was demonstrated by ten Lohuis and Miller (ten Lohuis and Miller, 1998), but comparisons to the *in hospite* condition have not been performed. Our results suggest that chromatin remodeling may play an important role in the switch between the free-living state and endosymbiosis, contributing to the substantial downregulation associated with the symbiotic life phase. We consider this resulting transcriptome reduction analogous to the genome reduction observed in numerous obligate symbiotic systems.

### The potential for sex in free-living Symbiodiniaceae

In many species, sex and reproduction are inextricably linked, but for some, sex represents a special occasion. For instance, consider a member of a taxon sister to the dinoflagellates, the apicomplexan *Plasmodium falciparum*. This malaria-causing parasite only engages in sexual reproduction within its mosquito host (Kooij and Matuschewski, 2007; Talman et al., 2004). Be it frequent or rare, the function of sex, despite its apparent ubiquity, is not a settled case. One hypothesis regarding the evolution of sex is that recombination mediated by sexual reproduction arose as a mechanism to repair DNA damage in an individual (Bernstein et al., 1985). A competing hypothesis holds that the advantage of meiotic recombination resides in the genetic diversity afforded to offspring (Burt, 2000; Otto, 2009).

The role of sexual reproduction in the Symbiodiniaceae life cycle remains an open question. A broad survey of Symbiodiniaceae by Santos and Coffroth (Santos and Coffroth, 2003) concluded that Symbiodiniaceae are diploid in the vegetative life phase, both in cultured and *in hospite* cells. However, apparent tetrads have long been reported in cultured Symbiodiniaceae (Freudenthal, 1962), but it remains unknown whether these tetrad-like arrangements represent meiotic events or simply two rounds of mitosis. Symbiodiniaceae meiosis has yet to be demonstrated in an experimental setting, but others have identified population genetic evidence of recombination (Baillie et al., 2000; LaJeunesse, 2001). Further, Symbiodinaceae genomes possess a virtually complete gene set for meiosis (Liu et al., 2018), suggesting that they have the potential for sexual reproduction.

We posit that the free-living phase of life in Symbiodinacaea may provide the opportunity for sex and recombination in the free-living state. In comparisons at 28°C, several meiosis-related transcripts are upregulated in the free-living state in comparison to the *in hospite* condition (*MSH6, RAD54*, *MEI2-like 2*, *MEI2-like 3*, and *MEI2-like 5*). Our work also identifies a subset of these meiotic transcripts (three *MEI2-like 5* transcripts) in the core heat stress response of *D. trenchii* that is shared by both *in hospite* and free-living states. While Levin et al. (2016) also detected an upregulation of meiosis-related genes under conditions of thermal stress in culture conditions, our work adds a nuanced understanding of alterations in meiosis-related gene expression between free-living and *in hospite D. trenchii*.

Recombination through sexual reproduction may be critical to the maintenance of Symbiodiniaceae genomes. Muller’s Ratchet dictates that asexual reproduction without other forms of recombination has the potential for the accumulation of deleterious mutations over generations (Felsenstein, 1974; Muller, 1964, 1932). This effect has support from RNA viruses lacking recombination (Chao, 1997) and may be one of the driving forces behind genome reduction in obligate endosymbionts (Andersson and Kurland, 1998), but obligate symbionts may have other mechanisms to avert genomic meltdown. Selection at the level of the holobiont has been suggested as one mechanism of endosymbiont genome maintenance (Rispe and Moran, 2000), with genomic degradation and losses observed likely to be neutral or nearly neutral (Pettersson and Berg, 2007). We posit that recombination in the free-living phase of Symbiodiniaceae may be instrumental in genomic maintenance of functions necessary for the free-living state.

### Evasion of host immunity by symbiont-derived factors

In order to maintain a stable endosymbiotic existence, members of the family Symbiodiniaceae must evade the host immune system. A considerable number of studies have shown that Symbiodiniaceae suppress and/or circumvent host immunity in order to colonize and persist *in hospite*; multiple mechanisms of circumvention appear to occur, including the suppression of apoptosis (Matthews et al., 2017; Rodriguez-Lanetty et al., 2006; Tchernov et al., 2011) as well as the manipulation of markers of endosome age (Chen et al., 2005, 2004, 2003). More recent work has identified endosymbiont miRNAs which target host transcripts involved in immunity (Baumgarten et al., 2018, 2013). We identified numerous transcripts involved in the suppression of innate immunity and anti-inflammatory pathways, including a putative NRCL3 ortholog and NF-κB inhibitor, which are upregulated in the symbiotic state. However, in order for this to plausibly occur, the symbiont must be able transmit these signals to the host. Based on transcriptome evidence, we postulate mechanisms for this below.

Apicomplexans, including parasites in the genus *Plasmodium,* possess structures that form an apical complex which functions in host invasion and secretes effectors into the host (reviewed by (Kemp et al., 2013). Cellular machinery in Symbiodinaceae involved in the delivery of immune-suppressing factors to the host remain unexplored, but this work identifies several abundant transcripts with homology to components of the apicomplexan apical complex. One component of the apical complex are micronemes, bar-like organelles which release micronemal proteins critical for host cell recognition and attachment, linking parasite and host cell receptors (Brecht et al., 2001; Carruthers and Tomley, 2008; Friedrich et al., 2010; Soldati et al., 2001) and further are critical for the trafficking and storage of ligands for host-cell receptors (Soldati et al., 2001).

*D. trenchii* possess numerous transcripts with homology to proteins associated with the apicomplexan apical complex, including three apical complex lysine methyltransferase transcripts (TRINITY_DN22464_c0_g1_i2, TRINITY_DN12687_c0_g1_i1, and TRINITY_DN59290_c0_g1_i1) and AGC/AKT protein kinase transcripts (TRINITY_DN23213_c0_g1_i1 and TRINITY_DN55685_c0_g1_i1), and three putative microneme proteins (TRINITY_DN25568_c0_g1_i1, TRINITY_DN48019_c0_g1_i1, and TRINITY_DN29950_c0_g1_i3). While the majority of these transcripts are not differentially expressed, two are highly upregulated *in hospite*. One of these transcripts (TRINITY_DN29950_c0_g1_i3) is a putative ortholog of a *Toxoplasma* microneme protein 4 (MIC4). In its role in infection, the *Toxoplasma* MIC4 is thought to contribute to parasite adhesion by acting as a bridge between receptors on the parasite and the host cell (Brecht et al., 2001). Another transcript upregulated *in hospite* with homology to apicomplexan invasion machinery is a cGMP-dependent protein kinase (TRINITY_DN29950_c0_g1_i3) which Collins et al. (Collins et al., 2013) show is required for the release of micronemes in *Plasmodium*. Taken together, these transcripts suggest the possibility of components of an apicomplexan-like apical complex involved in the maintenance of intracellular symbiosis by *D. trenchii*, and potentially the delivery of symbiont-derived factors which suppress the host immune response to allow stable endosymbiosis.

### Enhanced ammonium and ABC transporter activity *in hospite*

The controlled exchange of materials between the symbiont and host is of critical importance for the persistence of both partners (Behie and Bidochka, 2014; Moya et al., 2008). Our study identified transporters active in the symbiotic state for both ammonium transport and assimilation, as well as a diverse group of ABC transporters which may be involved in the movement of numerous as-yet unidentified substrates into and out the symbiont.

The acquisition of inorganic nitrogen is of critical importance to photosynthetic endosymbionts. Pernice et al. (Pernice et al., 2012) demonstrated that *in hospite* dinoflagellates fix 14 to 23 times more nitrogen than coral host cells in ammonium-enriched water. In the *in hospite* state, we identify the upregulation of an ammonium transporter transcript as well as two glutamine synthetase transcripts. This is congruent with the longstanding support for glutamine synthetase activity as the mechanism by which Symbiodinacaea assimilate ammonium from seawater and re-assimilate waste ammonium from the cnidarian host (Anderson and Burris, 1987).

In the *in hospite* state, five transcripts annotated as ABC transporters are upregulated in comparison to the free-living state at 28°C (Table S5). While numerous host ABC transporters are upregulated in the proteomes of symbiotic *Exaiptasia pallida* by Matthews et al. (2017), with earlier work by Peng et al. (2010) identifying an ABC transporter associated with the symbiosome, this represents the first functional evidence *in hospite* Symbiodiniaceae transporter activity in a cnidarian. ABC transporters comprise a family of proteins responsible for the transport of diverse substrates, including ions, nutrients, amino acids, peptides, proteins, lipids, metals, lipids, oligonucleotides, and sugars (Locher, 2016; Wilkens, 2015).

### Conclusion

In summary, our results demonstrate that symbiotic state has major impacts on Symbiodiniaceae gene expression, both in steady state symbiosis and under conditions of acute thermal stress. While prior work has found physiological differences between *in hospite* and free-living Symbiodiniaceae (e.g. Banaszak and Trench, 1995; Bhagooli et al., 2008), effect of the intracellular habitat on the transcriptome has not previously been appreciated. The *in hospite* response to thermal stress differs dramatically from that of free-living *D. trenchii*, with the transcriptome of *in hospite* endosymbionts hallmarked by higher differential gene expression activity overall and enriched for mRNA transport activity as well as a specialized cell adhesion process. While we identify a core heat stress response shared between free-living and *in hospite D. trenchii*, this represents only a minority of transcriptional activity. Our findings also reveal the divergent processes at work in free-living and *in hospite* cells under non-stress conditions, with free-living *D. trenchii* exhibiting evidence suggestive of sexual reproduction and *in hospite* cells displaying reduced differential expression in a process we term transcriptome reduction, as well as putative host evasion activity. We posit that the observed transcriptomic reduction of non-essential genes *in hospite* may be an alternative adaptive strategy to genome reduction for endosymbionts that must also maintain the capacity for a free-living life phase. We intend for this to serves as the basis for future gene editing efforts to understand the genetic basis of symbiosis and dysbiosis, with the ultimate aim of engineering of stress-tolerant coral endosymbionts. These efforts will for provide additional options in future reef protection and restoration efforts desperately needed for the persistence of coral reefs into the 21^st^ century.

## Materials and Methods

Aposymbiotic *Exaiptasia pallida* (strain CC7) were inoculated with *D. trenchii* (strain CCMP2556) and maintained with feeding of *Artemia franciscana* twice weekly for 12 months. Free-living *D. trenchii* were cultured in Prov50 (Smith and Chanley, 1975) medium and maintained in log phase via regular passage. To identify thermal treatments that elicited an acute heat stress response, *Durusdinium trenchii* (strain CCMP2556) cultures were exposed to thermal stress and assessed with PAM fluorometry (Fig. S1). Based upon the observed rapid decline in Fv/Fm at 34 °C in contrast to the maintained photosynthetic efficiency at 28 °C (Fig. S1), these treatments were selected and implemented for comparisons following 6 hours at the target temperatures. Symbiotic anemones and free-living *D. trenchii* cultures were held at 28°C under light intensity levels of 80 μE (12:12 light:dark cycle) and ramped up to the 34°C challenge at a rate of 2°C per day for thermal stress treatments, with increases of temperature performed at the beginning of the photoperiod. Free-living (cultured) *D. trenchii* and *in hospite D. trenchii* were frozen in liquid nitrogen mid-photoperiod for sampling. Heat stress treatments were exposed to 34°C for a total of six hours. The experiment was replicated temporally (n=3-4). Total RNA was extracted from Symbiodiniaceae cells, with *in hospite* samples depleted of cnidarian host material prior to extraction using a novel endosymbiont enrichment method (See Supplement 1 for detailed methods). Stranded RNAseq libraries were prepared from poly(A) selected RNA with the NEBNext Ultra RNA Library Prep Kit (NEB, Cambridge, MA) and sequenced across two Illumina HiSeq lanes. A reference transcriptome was assembled and filtered to remove host, other non-Symbiodiniaceae transcripts, and ribosomal RNA transcripts. Principal component analysis was conducted using the prcomp function from the stats package within R and statistical inference of significant transcripts driving the principal components was conducting using the jackstraw package in R. The resulting transcriptome was annotated using Trinotate (https://trinotate.github.io/) with BLASTX and BLASTP searches performed against the SwissProt and Uniref90 databases using DIAMOND, an accelerated sequence aligner comparable to NCBI BLAST+ (Buchfink et al., 2015). Following Trinotate, we employed the structural protein homology detection program HH-Pred and the PDB70 database to identify remote homologs in all remaining un-annotated transcripts using a strict cut-off of 80% probability. Annotation was further supplemented by using Blast2GO (Conesa et al., 2005) in combination with BLASTX searches against the non-redundant NCBI protein database with DIAMOND and Interproscan under default settings. Gene ontology terms acquired for each transcript during the annotation process from Trinotate, HHPred (Söding et al., 2005), and Blast2GO were merged and redundant terms resulting from the integration of the three annotation tools were removed prior to downstream analyses. Following transcript quantification with Salmon (Patro et al., 2017), differential gene expression analysis was performed in edgeR (Robinson et al., 2010) using a generalized linear model to determine the effects of temperature and symbiotic state as well as the interaction between the two on changes in transcript expression. Differentially expressed transcripts were subject to gene ontology (GO) enrichment analyses within the R package topGO (Alexa and Rahnenführer, 2016). GO results for comparisons were visualized using REVIGO (Supek et al., 2011). Methods are described in detail in Supplement 1.

## Supporting information

Supplement_2_DEGs_Annotations.xlsx

## Supplement 1

### Supplemental Materials and Methods

#### Experimental Overview

Briefly, the experimental design compares D. trenchii in the free-living state (culture) and in hospite in an Exaiptasia host under two different temperature regimens, a steady-state culture temperature of 28°C and a thermal stress of 34°C, both under light intensity levels of 80 μE. Four biological replicates of each condition were implemented, with replication performed in time.

#### Exaiptasia symbiont manipulation and Symbiodinacaea culture

Aposymbiotic Exaiptasia were produced through repeated cold treatment (8 hours at 4°C, repeated three times over the course of one week) followed by DCMU treatment (50 uM) performed for 6 hours with 1000 μmol m−2 s−1 light intensity, repeated twice weekly for 3 weeks. Absence of dinoflagellates was confirmed with fluorescent microscopy. D. trenchii strain CCMP2556 was obtained from the National Center for Marine Algae and Microbiota (East Boothbay, Maine) and cultured in Prov50 media under a 12:12 light cycle. Log phase cultures were used for all experiments. Aposymbiotic anemones were challenged with 100,000 log phase Durusdinium trenchii cells combined with freshly hatched Artemia franciscana in a total volume of 10 ml of sterile seawater, with water replaced after 12 hours. Infection was confirmed by fluorescent microscopy, followed by PCR and sequencing of 23S. Anemones hosting D. trenchii were grown for one year prior to experimental manipulation with light levels at approximately 80 μmol m−2 s−1. Anemones were fed freshly hatched Artemia twice weekly.

#### Symbiosis state comparison and thermal challenge

To inform the design of this experiment, maximum quantum yield in cultured D. trenchii was assessed through time, at temperatures from 28 to 36° C (n=3) using a PAM fluorometer (Walz, Germany). We found that the rapid decline in maximum quantum yield began at 34°C at 2 hours of thermal stress. In an 8 hour timespan, we did not find significant decline at the lower thermal treatments.

Anemones and cultures were maintained at 28°C with 80 μmol m−2 s−1 irradiance and ramped up to the 34° C challenge at a rate of 2°C per day for thermal stress treatments. Mid-photoperiod (6 hours into light phase of the diel cycle), samples were frozen for subsequent RNA extraction. Log phase D. trenchii cultures were used for all experiments. Replicates were performed in time, with one replicate of each treatment included in each trial. A total of four biological replicates were collected for each treatment.

#### Symbiodinacaea RNA Extraction Enrichment Protocol

Anemones were frozen in minimal seawater in 2 ml cryotubes in liquid nitrogen and stored at −80°C till extraction. For enrichment of Symbiodiniaceae from anemone extractions, 500 ul of Buffer RLT with 0.1% 2-mercaptoethanol (Qiagen) was added to each tube and manually flicked till frozen tissue was dislodged and decanted into 1.5 ml centrifuge tube. Tissue was gently homogenized with a plastic pestle and vortexed at maximum speed for 1 minute. Homogenate was centrifuged at 10,000 g for 3 min. Clear supernatant was carefully removed and discarded, leaving the symbiont pellet intact.

This pellet was resuspended in 1.4 ml of Buffer RLT and vortexed till the mixture was homogenous. This suspension was centrifuged at 3,000 g for 3 min. Clear supernatant was discarded, and an additional 1.4 ml of Buffer RLT was added vortexed till homogenous. This was centrifuged at 3,000 g for 3 minutes and clear supernatant was discarded prior to resuspension in 600 ul Buffer RLT. Contents were transferred to a screw top 2.0 ml tube and bead beat for 1 min at 6 m/s with glass beads. Homogenate was processed with the Qiagen RNeasy Kit as per manufacturer instructions, with an additional RPE Buffer wash prior to elution of total RNA. Symbiodiniaceae were snap frozen in liquid nitrogen and extracted alongside anemone tissue, implementing an identical washing scheme to eliminate any extraction-based bias.

#### Library Construction and Sequencing

Following poly(A) selection from total RNA, RNAseq libraries were constructed from mRNA using the NEBNext Ultra Directional RNA Library Prep Kit (NEB). A total of 16 libraries were sequenced on two Illumina HiSeq lanes, using v4 150 base pair paired-end chemistry (Illumina). The indexed samples were pooled across both lanes to eliminate lane effects. Raw data have been archived on the SRA and is available under NCBI TSA BioProject PRJNA508937.

#### De novo transcriptome assembly

Quality of the resulting raw reads were assessed via FastQC and low-quality reads were removed and/or trimmed with Trimmomatic (Bolger et al., 2014) with a minimum read length threshold of 50 bp. Trimmed reads that aligned concordantly against the human and Exaiptasia pallida genomes using the HISAT2 (Kim et al., 2015) program to remove any contaminating eukaryotic reads. Remaining reads were then filtered using all bacterial, archaeal, and viral genomes present in RefSeq (as of Sep. 8, 2017) using Kraken, an ultra-fast taxonomic sequence classification program (Wood and Salzberg, 2014). The 526,545,945 remaining PE reads were then assembled with Trinity v2.4.2 (Grabherr et al., 2011) with a minimum contig length of 200 bp.

Redundant contigs were removed using uCLUST (Edgar, 2010) at an identity threshold of 100%. Removal of eukaryotic contaminants via Hisat2 and prokaryotic contigs via Kraken (Wood and Salzberg, 2014) was repeated once more on the transcriptome assembly to remove any lingering contamination that could not be identified from the raw reads. Ribosomal RNA contamination was identified and removed by aligning transcriptome contigs against all of the SILVA rRNA databases (Quast et al., 2012) with Bowtie2 (Langmead and Salzberg, 2012) resulting in a final assembly of 85,130 contigs. Assembly statistics were calculated using Transrate (Smith-Unna et al., 2016) (Supplementary Table 1).

For annotation, the assembly was searched against the NR protein database using DIAMOND, a fast search algorithm that employs double-indexing and spaced seeds of various weights and shape to make it ~20,000x faster than BLASTX with comparable level of sensitivity and accuracy (Buchfink et al., 2015). These results were input into with Trinotate using BLASTP and BLASTX searches against the SwissProt and Uniref90 databases with an e-value cutoff of < 1 × 10^−5^. Open reading frames (ORFs) were predicted using Transdecoder under the default settings. Gene ontology classifications were extracted from search results against the SwissProt database. The searches were conducted using DIAMOND, a fast search algorithm that employs double-indexing and spaced seeds of various weights and shape to make it ~20,000x faster than BLAST with a similar sensitivity and accuracy (Buchfink et al., 2015). Additional annotation was performed using Blast2GO (Conesa et al., 2005), with redundant GO terms removed with a custom script.

Quality-controlled reads were mapped back to the final assembly using Salmon at an average mapping rate of 88% and 30.1 million reads per sample. Mapping results were imported into R for differential gene expression analysis with the edgeR package (Robinson et al., 2010). Low-count transcripts that across all samples with a mapping rate of < 10 reads were removed prior to count normalization within R. Differential gene expression anlysis was then performed in edgeR using a generalized linear model to determine the effects of temperature and symbiotic state as well as the interaction between the two on changes in gene expression. Four pairwise comparisons were then conducted using the GLM: (1) free-living at 28 °C vs. free-living at 34 °C (2) symbiotic at 28°C vs. symbiotic at 34 °C (3) free-living at 28 °C vs. symbiotic at 28 °C and (4) free-living at 34 °C and symbiotic at 34 °C. An additional comparison with the GLM was done using the formula ((symbiotic at 34 °C vs. free-living at 34 °C) - (free-living at 28 °C vs. symbiotic at 28 °C)) to determine genes that were differentially expressed in response to symbiosis but responded differently to temperature. Results from each of the five comparisons in edgeR were then used for gene ontology (GO) enrichment analyses within the R package topGO. GO results for the different comparisons were summarized and viewed using the web-based REVIGO program (Supek et al., 2011).

**Supplementary Figure 1.**
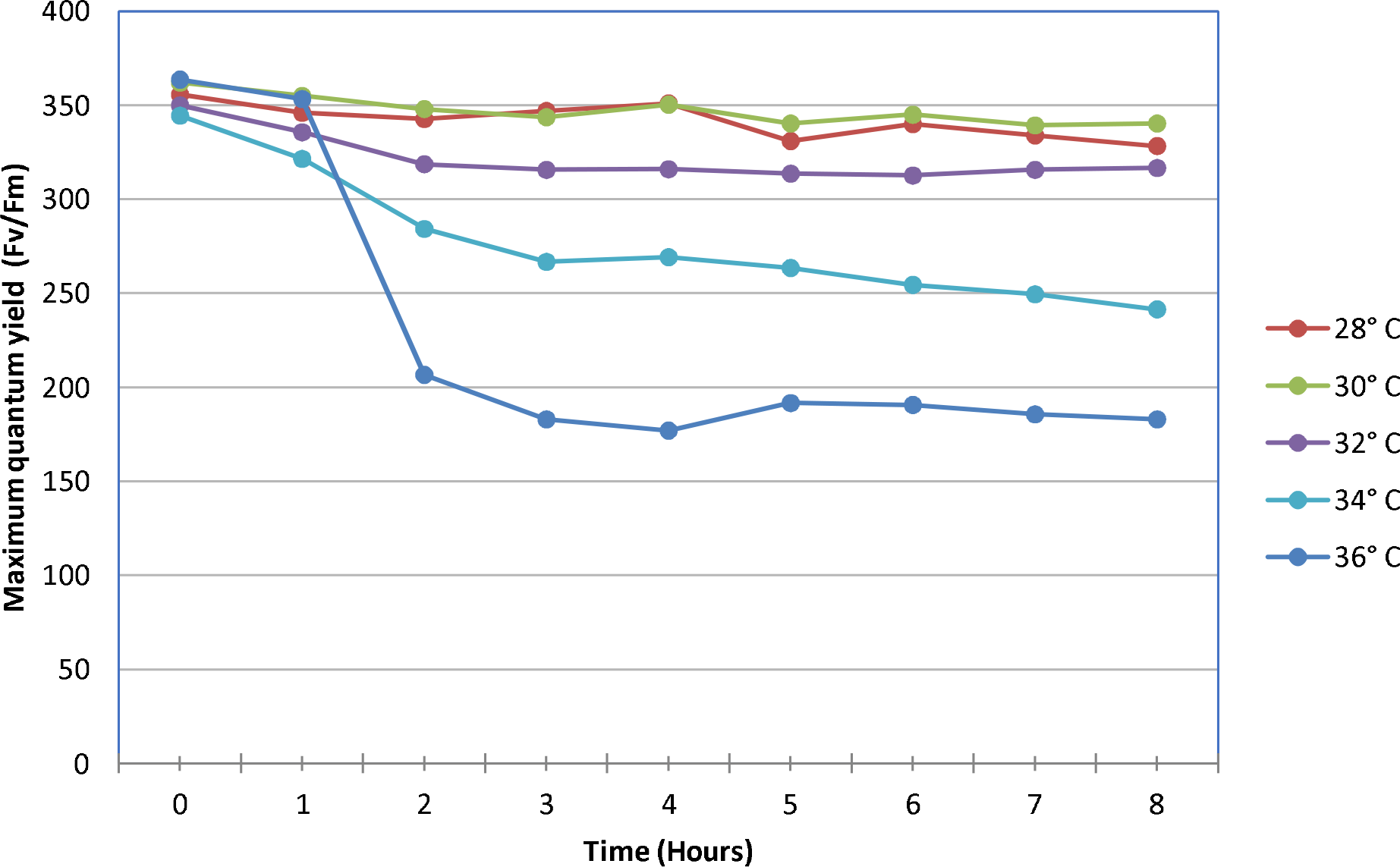
Maximum quantum yield of D. trenchii cultures assessed via PAM fluorometry throughout a thermal stress timecourse.

**Supplementary Figure 2.**
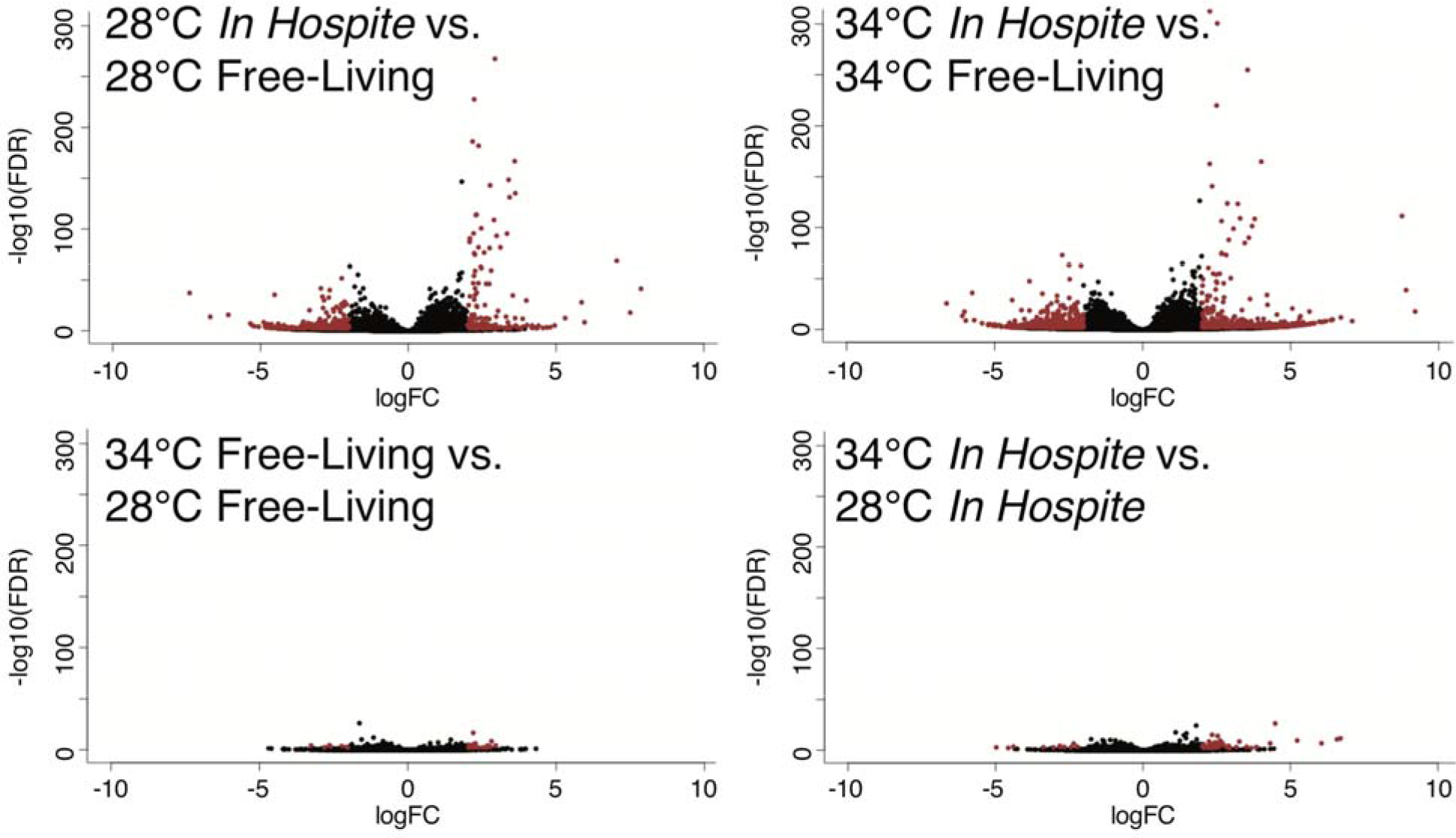
Volcano plots of all transcripts; transcripts with FDR corrected p-value < 0.01 and greater than log2 2-fold difference are indicated in red.

**Supplementary Figure 3.**
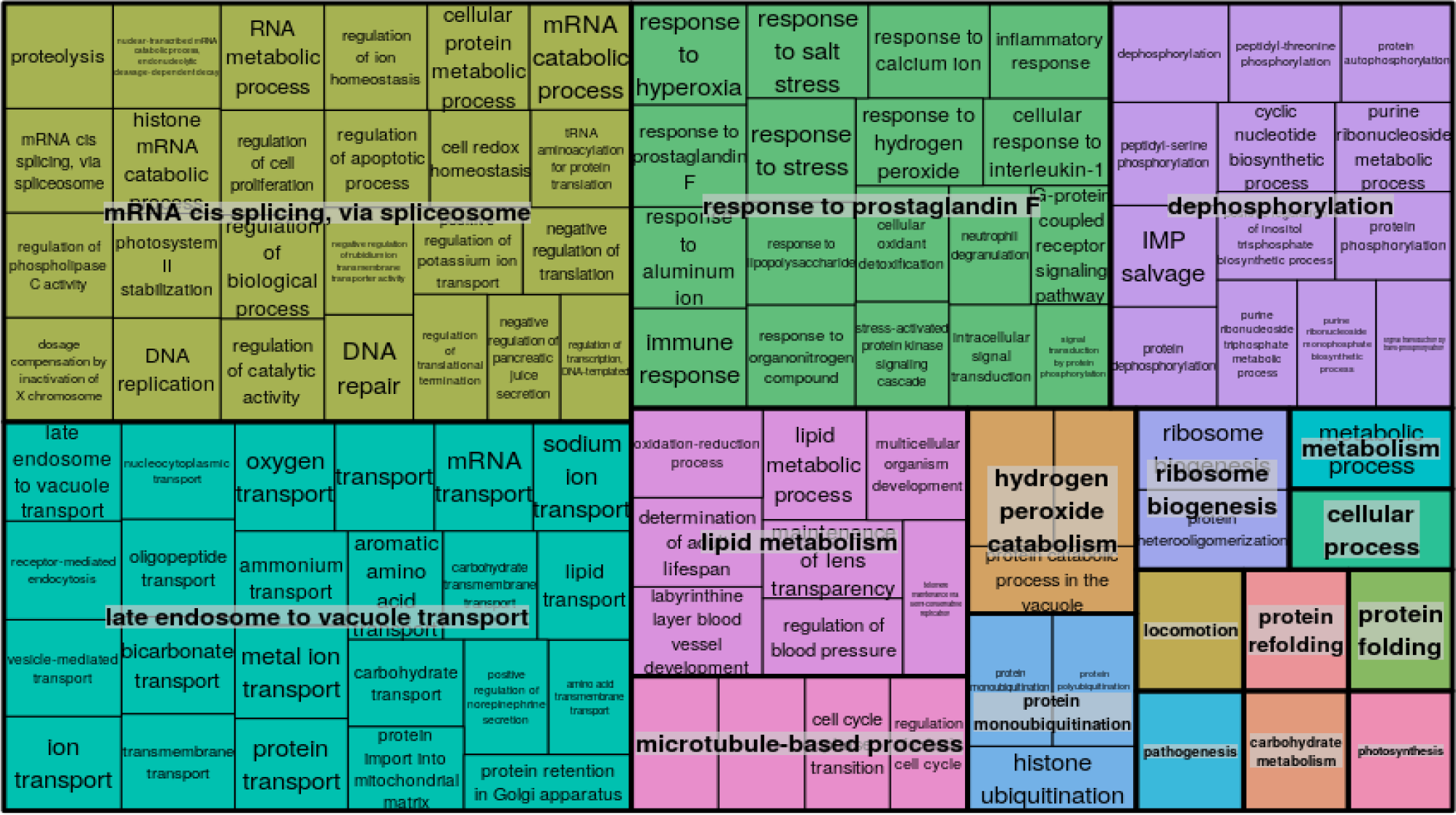
REVIGO tree diagram summary of the GO terms associated with transcripts up-regulated in response to symbiosis compared to free-living D. trenchii at 28 °C.

**Supplementary Figure 4.**
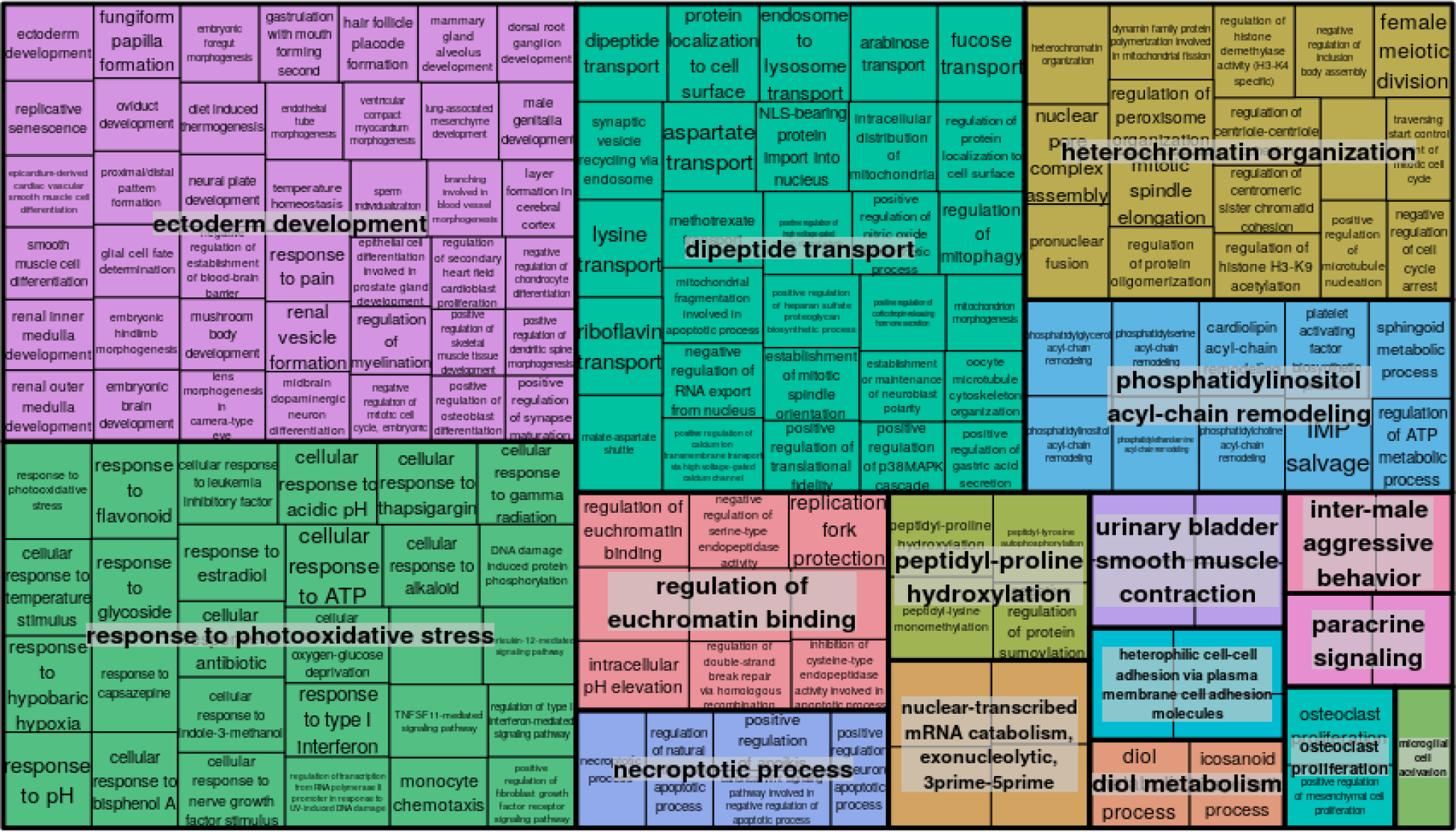
REVIGO tree diagram summary of the GO terms associated with transcripts down-regulated in response to symbiosis compared to free-living D. trenchii at 28 °C.

**Supplementary Figure 5.**
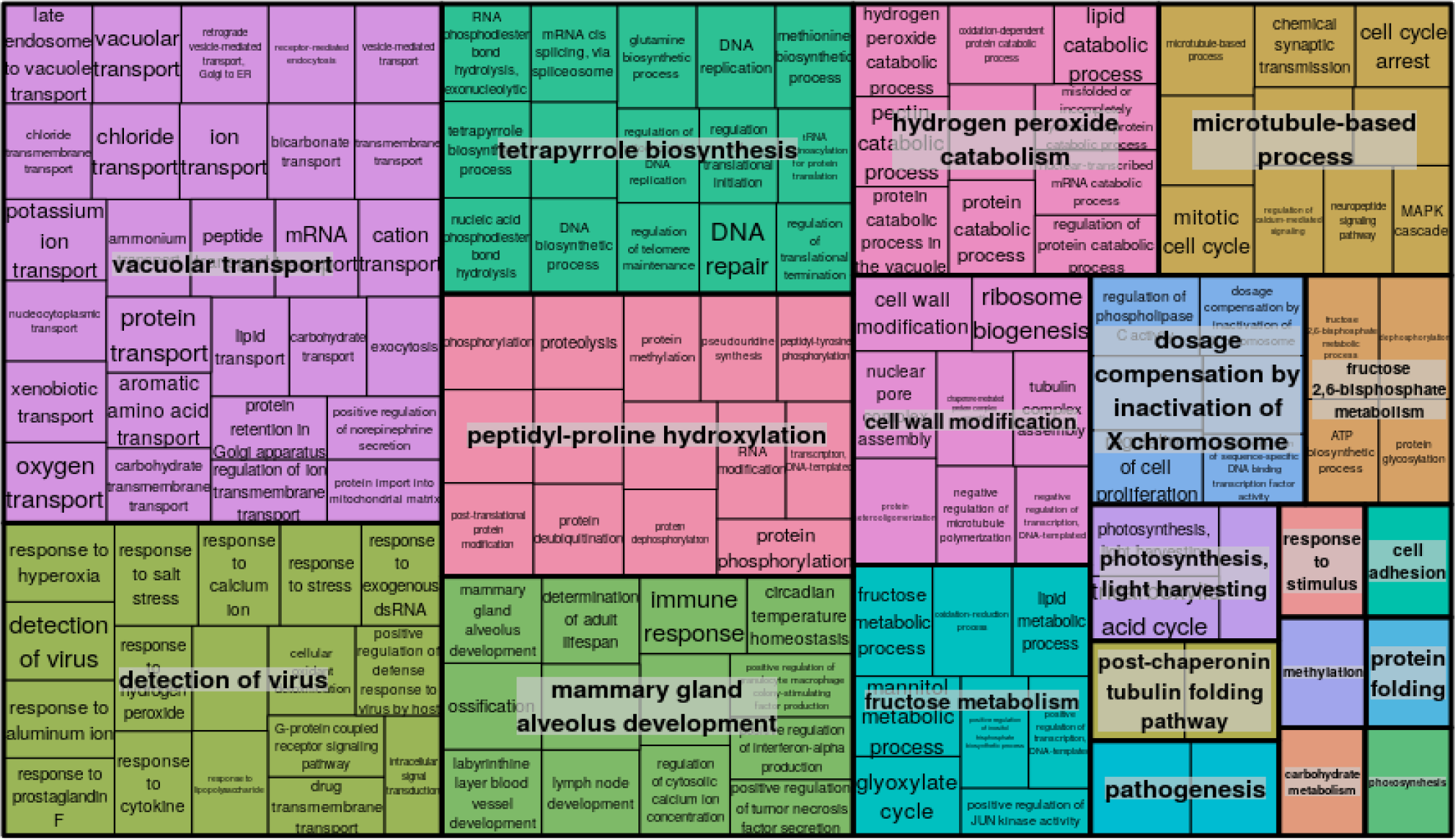
REVIGO tree diagram summary of the GO terms associated with transcripts up-regulated in response to symbiosis compared to free-living D. trenchii at 34 °C.

**Supplementary Figure 6.**
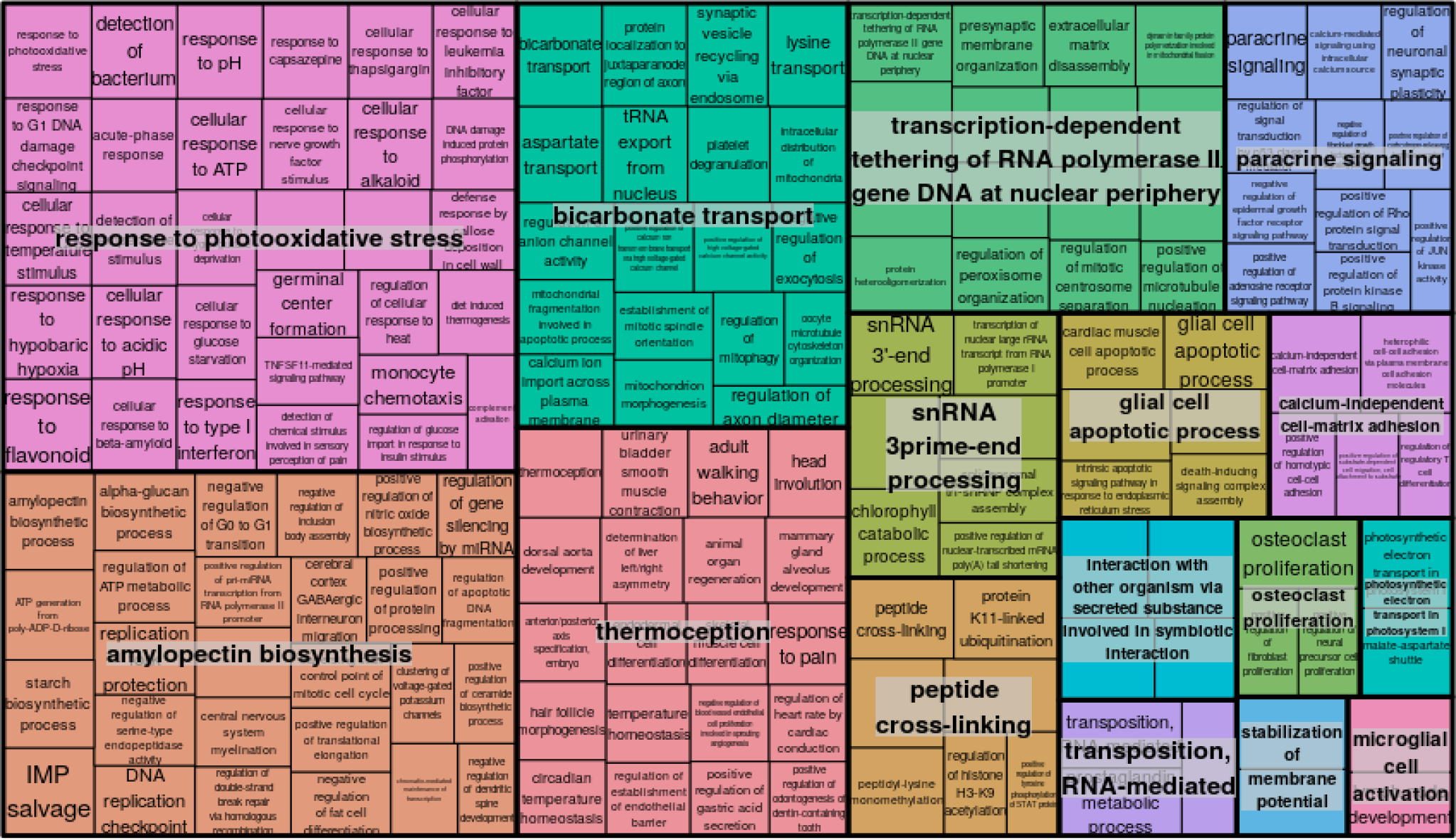
REVIGO tree diagram summary of the GO terms associated with transcripts down-regulated in response to symbiosis compared to free-living D. trenchii at 34 °C.

**Supplementary Figure 7.**
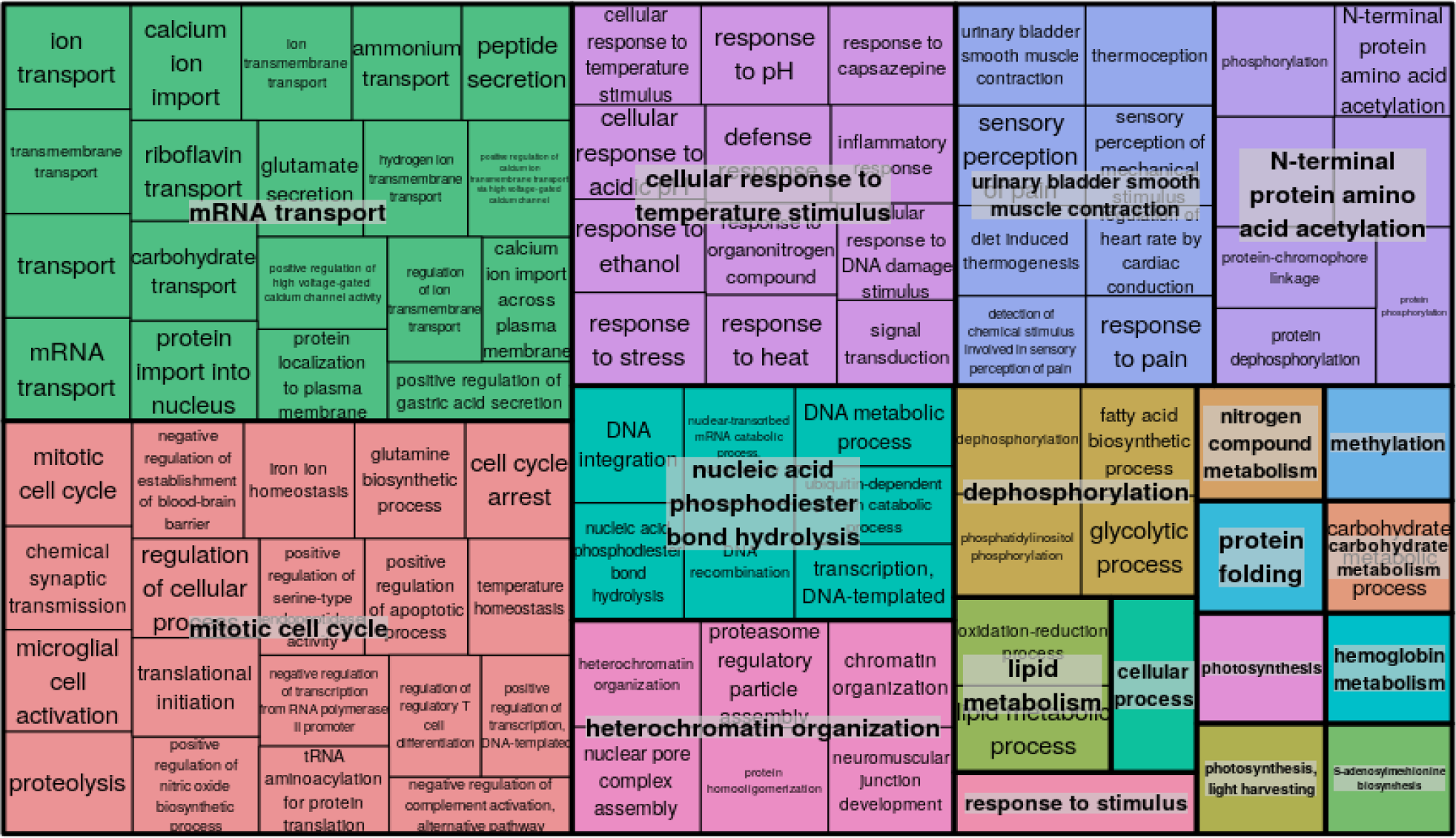
REVIGO tree diagram summary of the GO terms associated with transcripts up-regulated in response to temperature stress in in hospite D. trenchii at 34 °C.

**Supplementary Figure 8.**
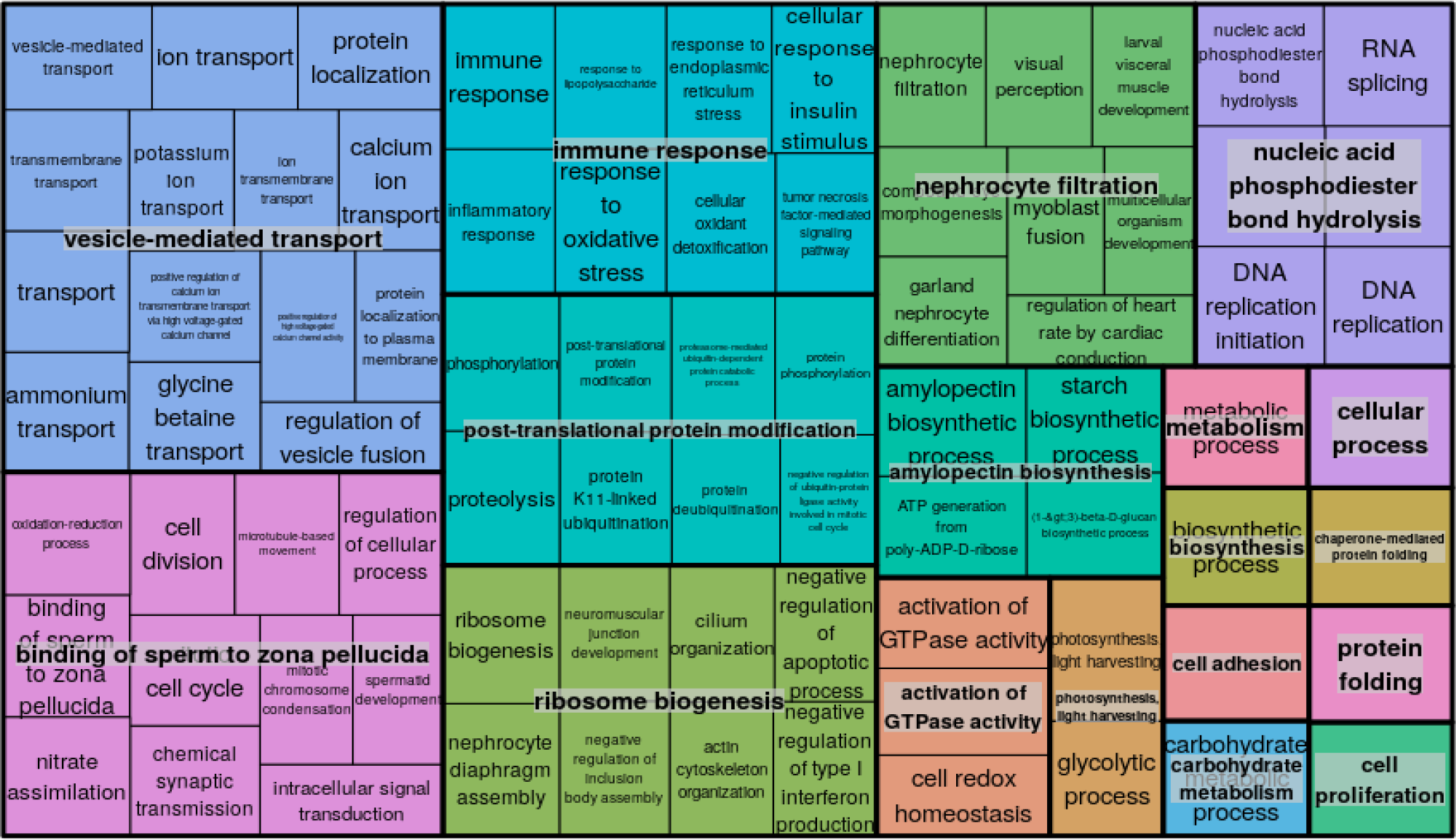
REVIGO tree diagram summary of the GO terms associated with transcripts down-regulated in response to temperature stress in in hospite D. trenchii at 34 °C.

**Supplementary Figure 9.**
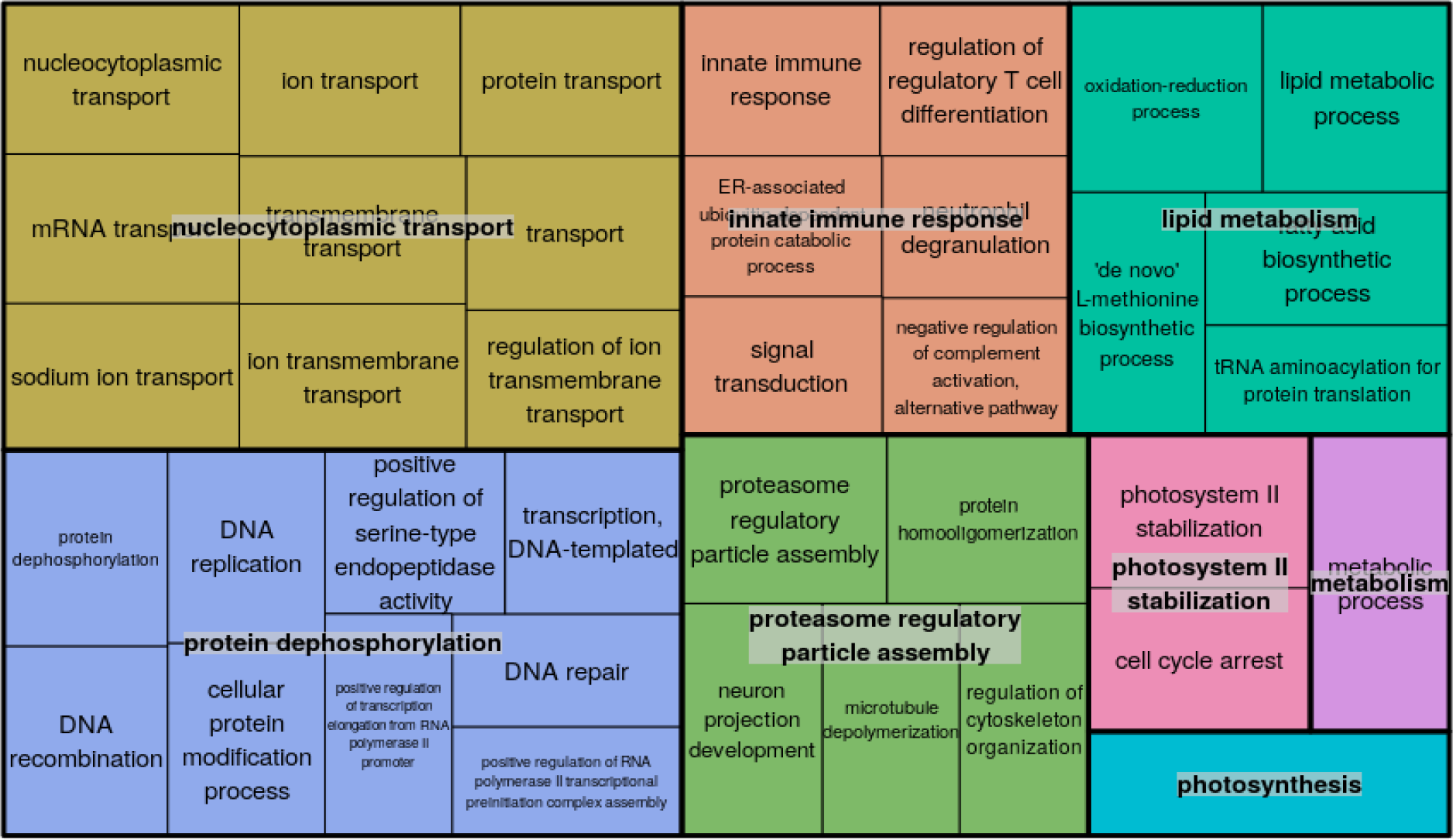
REVIGO tree diagram summary of the GO terms associated with transcripts up-regulated in response to temperature stress in free-living D. trenchii at 34 °C.

**Supplementary Figure 10.**
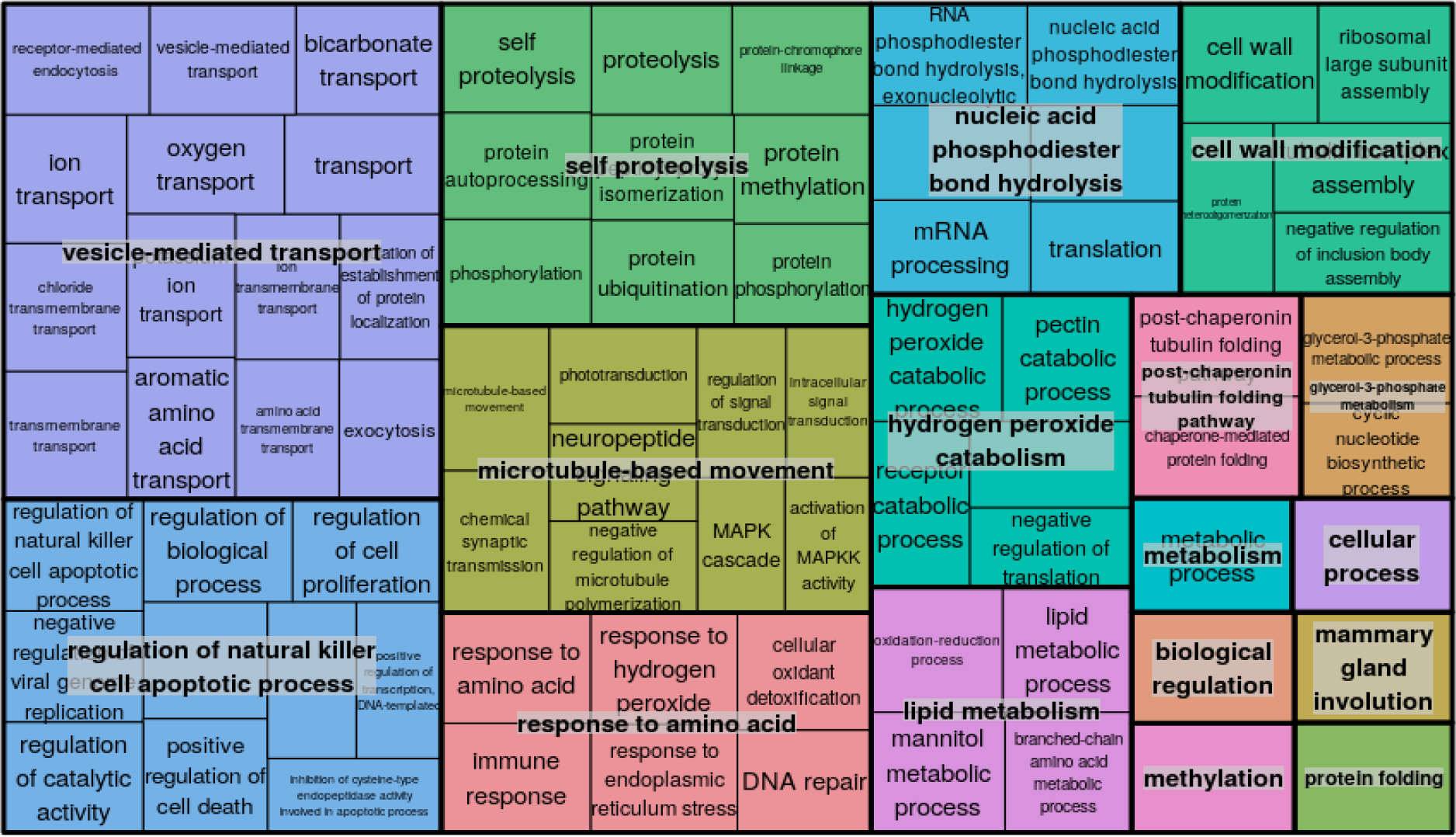
REVIGO tree diagram summary of the GO terms associated with transcripts down-regulated in response to temperature stress in free-living D. trenchii at 34 °C.

**Supplementary Figure 11.**
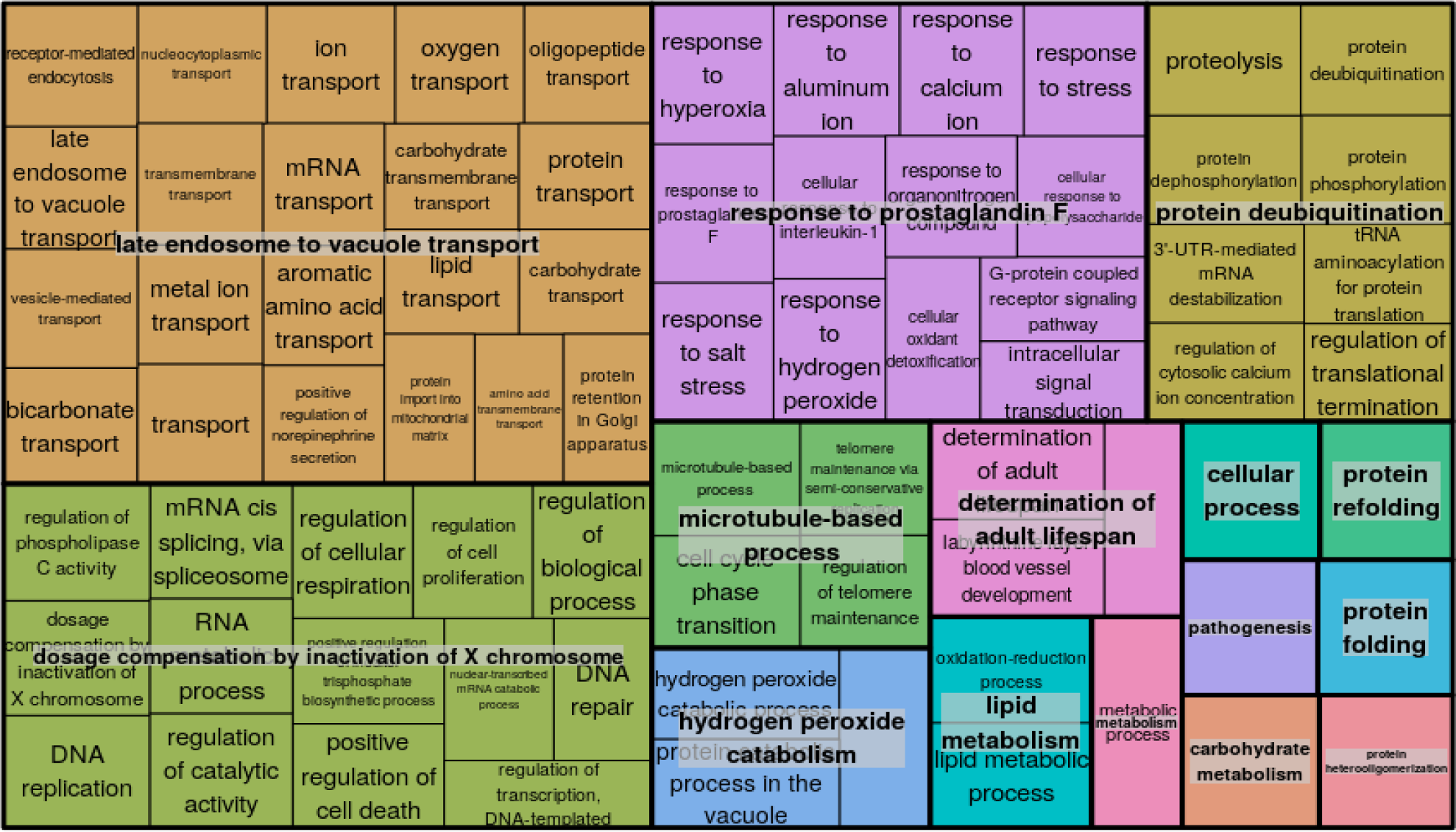
REVIGO tree diagram summary of the GO terms associated with transcripts that were up-regulated in response to symbiosis in D. trenchii at both 28 °C and 34 °C.

**Supplementary Figure 12.**
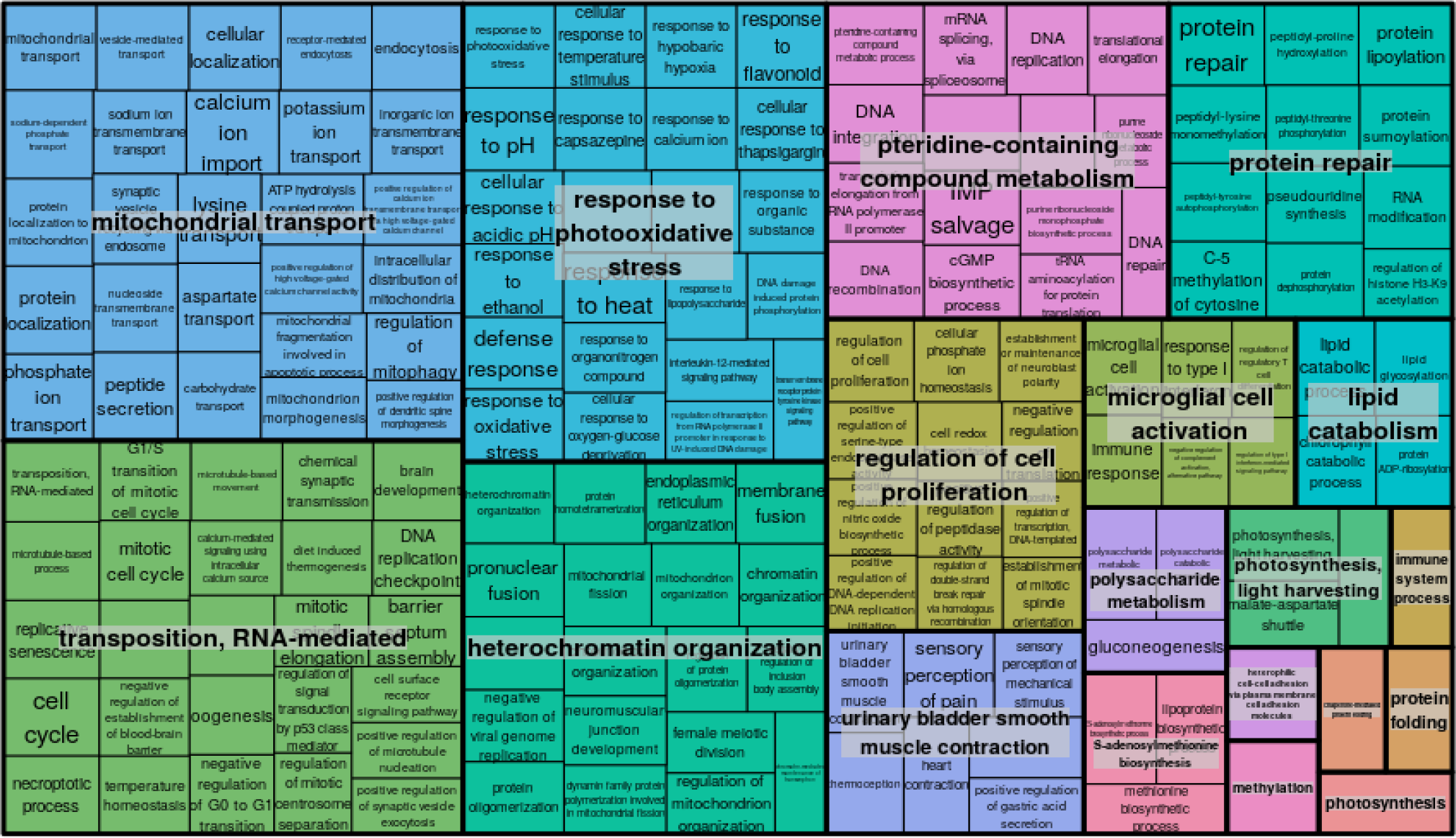
REVIGO tree diagram summary of the GO terms associated with transcripts that were down-regulated in response to symbiosis in D. trenchii at both 28 °C and 34 °C.

**Supplementary Figure 13.**
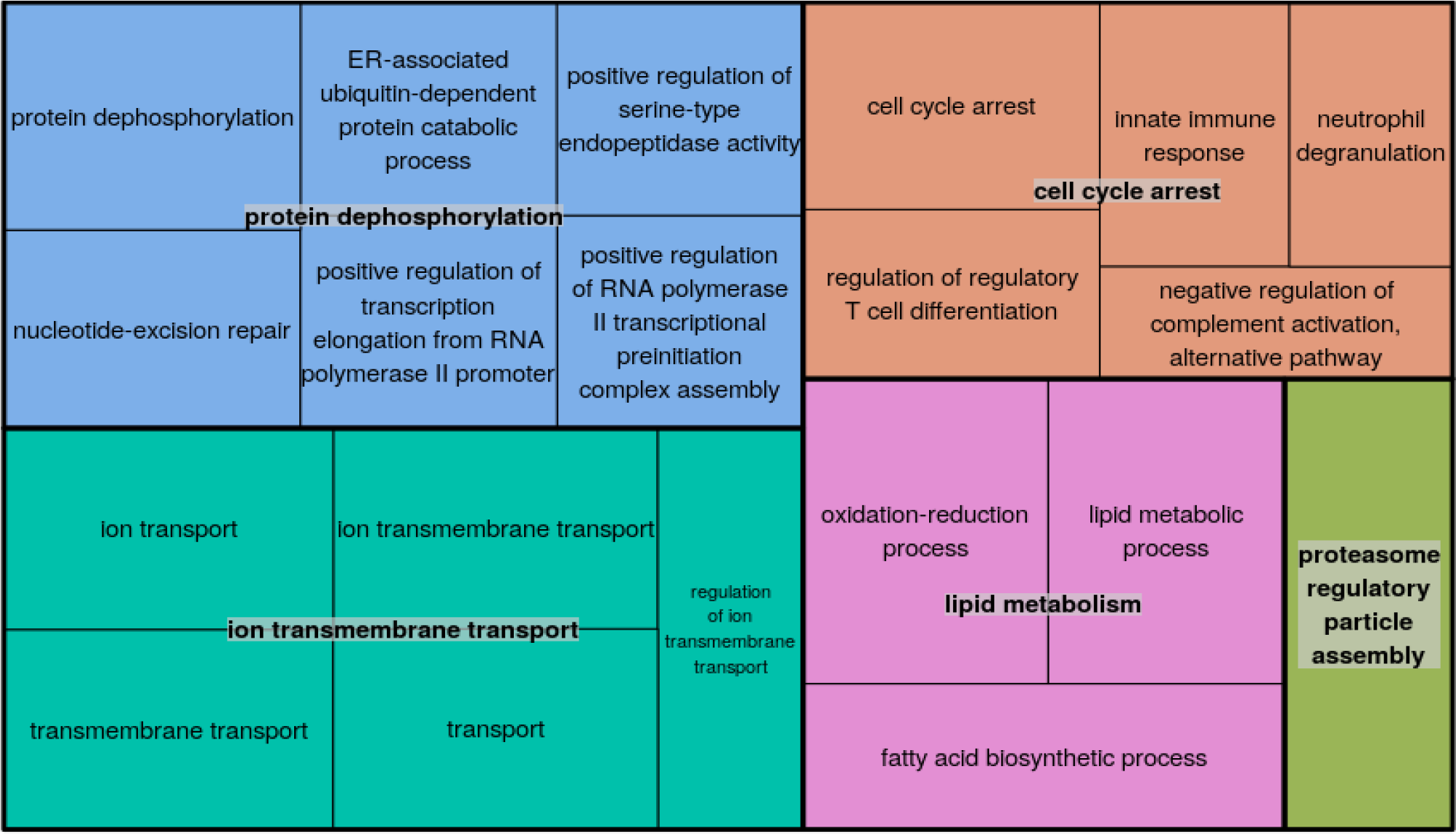
REVIGO tree diagram summary of the GO terms associated with transcripts that were up-regulated in response to temperature stress in both free-living and symbiotic D. trenchii.

**Supplementary Figure 14.**
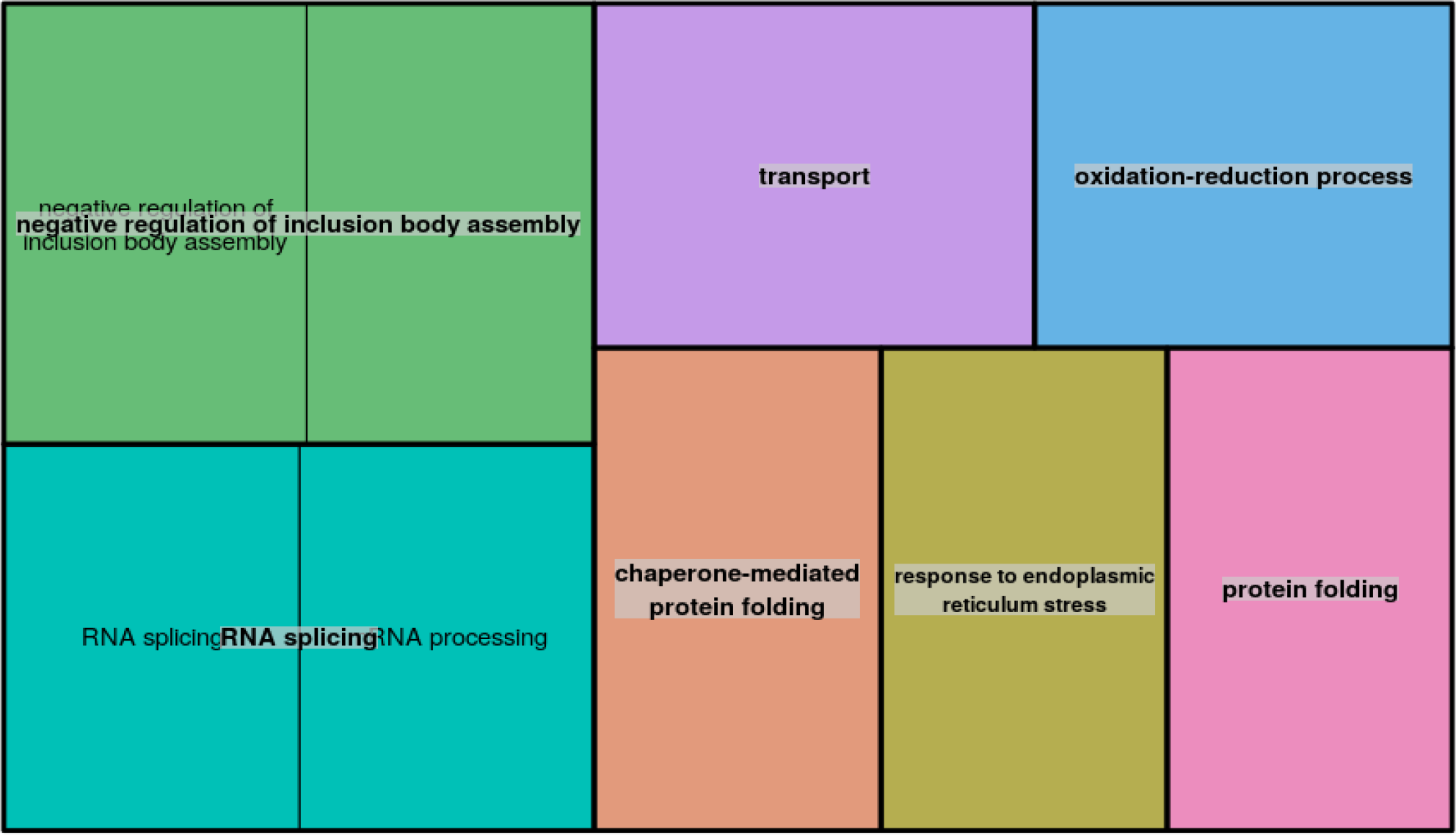
REVIGO tree diagram summary of the GO terms associated with transcripts that were down-regulated in response to temperature stress in both free-living and symbiotic D. trenchii.

**Supplementary Table 1.** Assembly statistics for the D. trenchii assembly

| **Assembly metrics** |  |
| --- | --- |
| **Total PE read sets after QC** | 526,545,945 |
| **Total PE read sets after QC (%)** | 81.5% |
| **Assembly length (Mbp)** | 81.7 |
| **# Transcripts** | 85,168 |
| **Predicted ORF count** | 47,335 |
| **Genes annotated** | 28,531 |
| **Genes annotated (%)** | 60.3% |
| **Longest gene length (bp)** | 8,088 |
| **Mean gene length (bp)** | 960.36 |
| **N50 (bp)** | 1,470 |
| **GC content (%)** | 0.55 |

**Supplementary Table 2.**
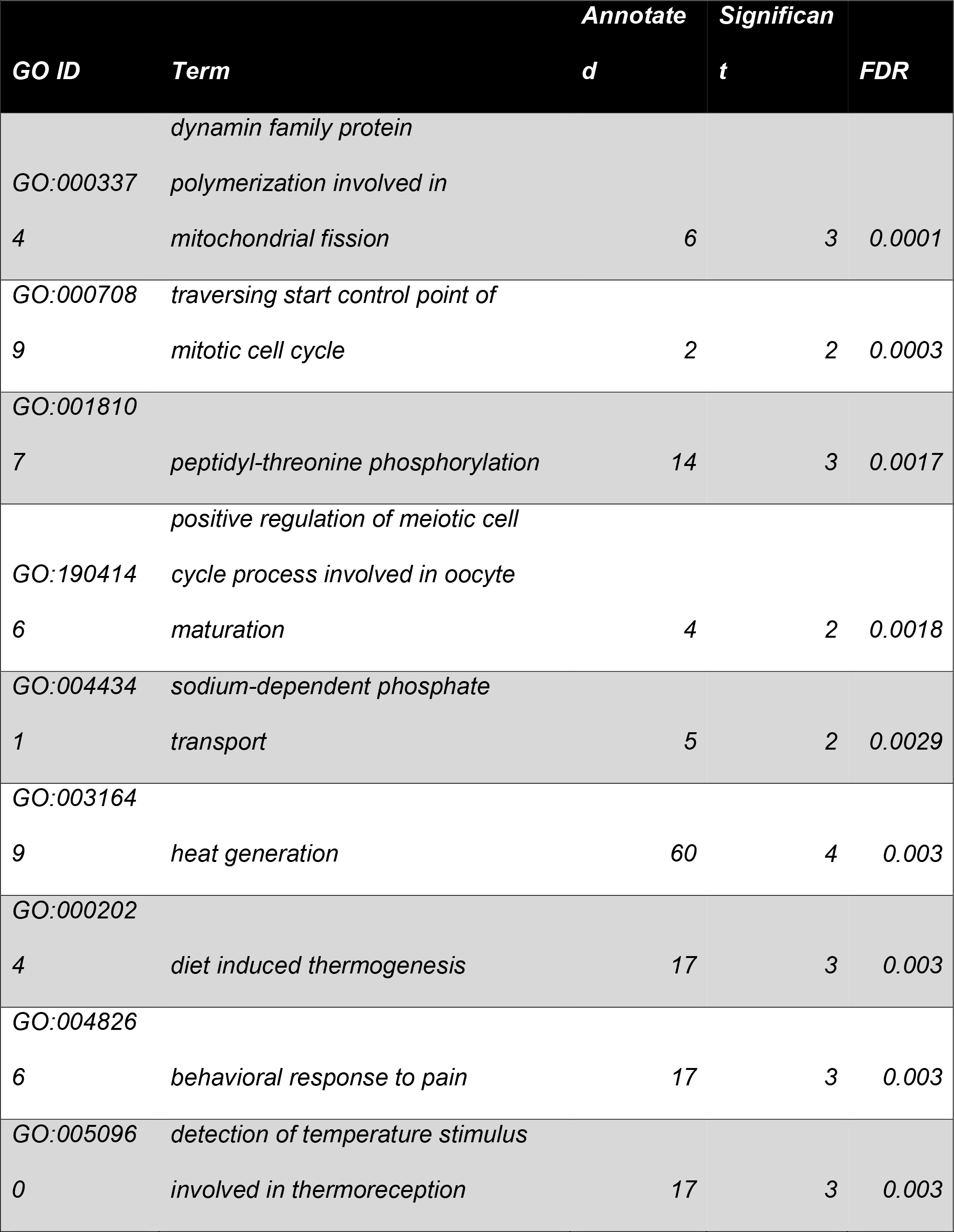
Gene ontology enrichment analysis results comparing the transcriptional profiles of D. trenchii when in hospite at 28 °C to the free-living stage at 28°C

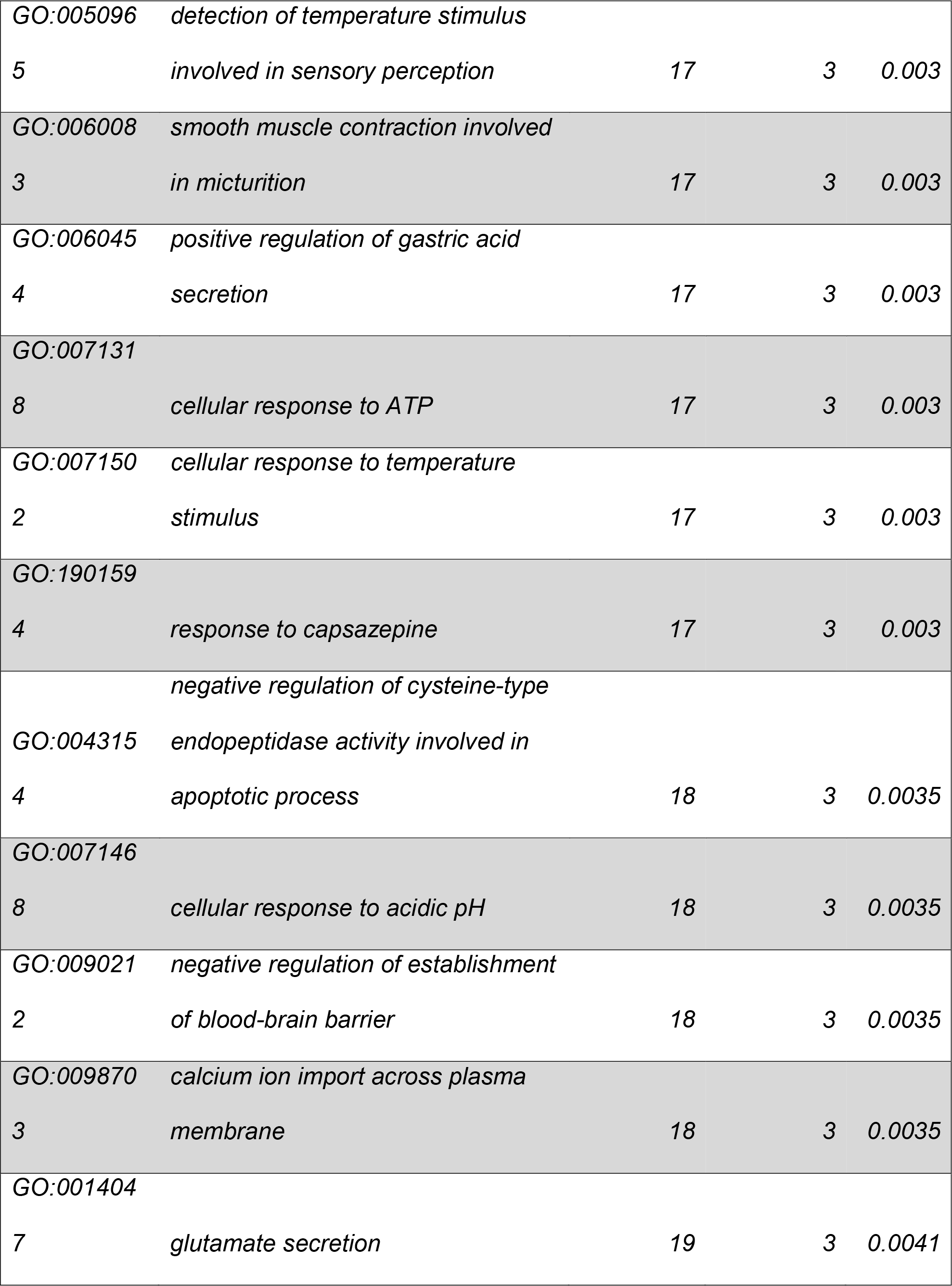

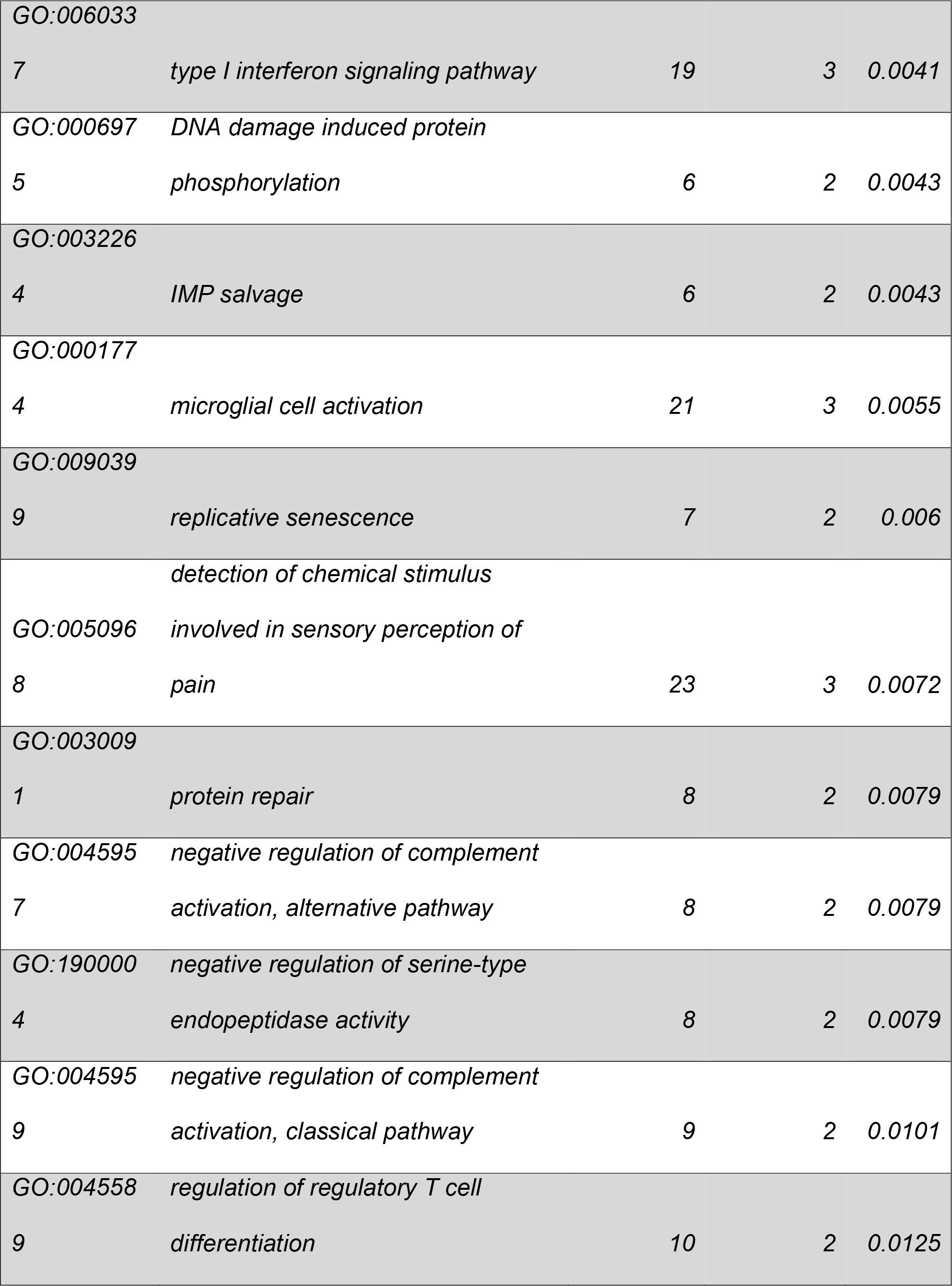

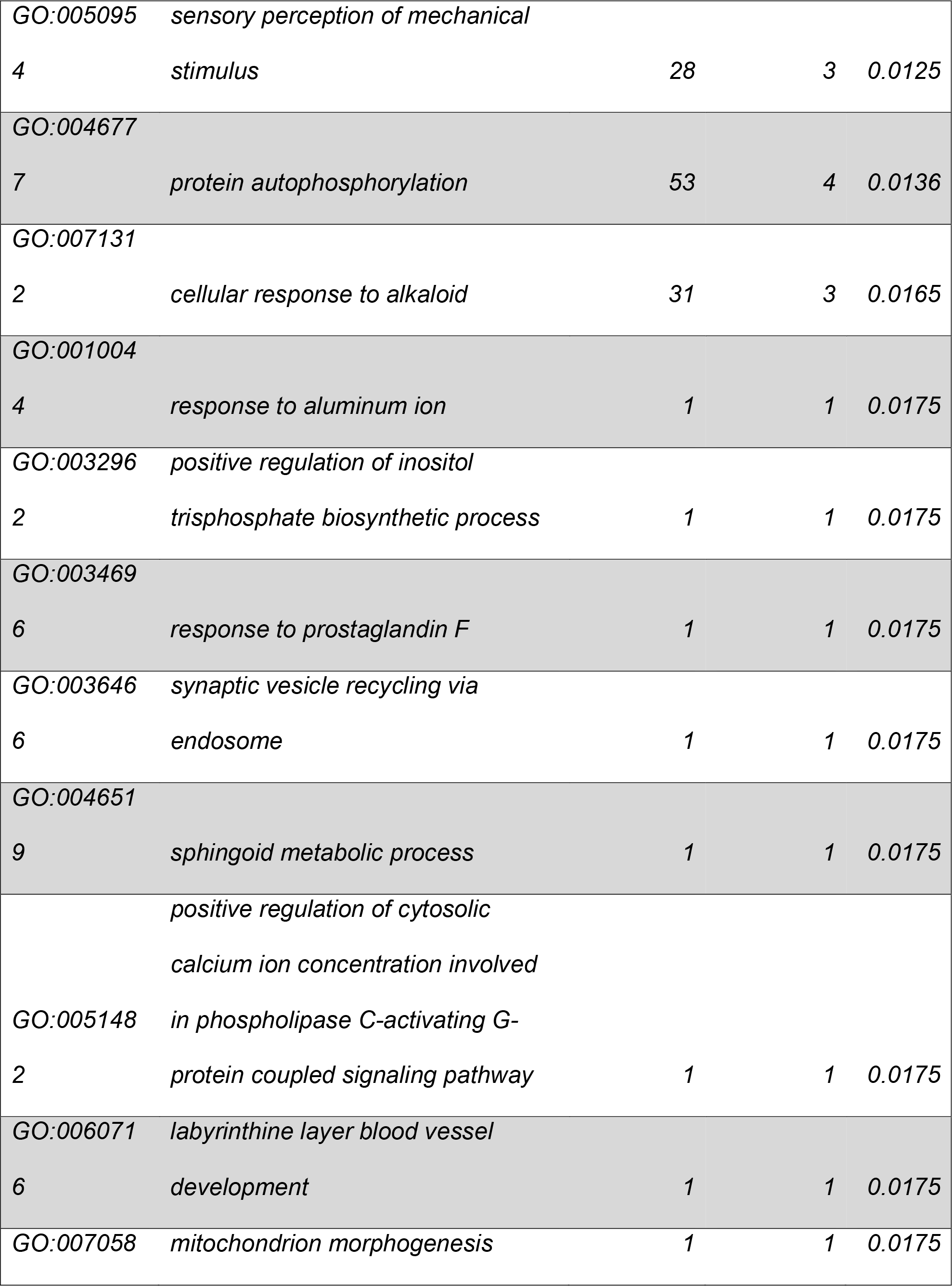

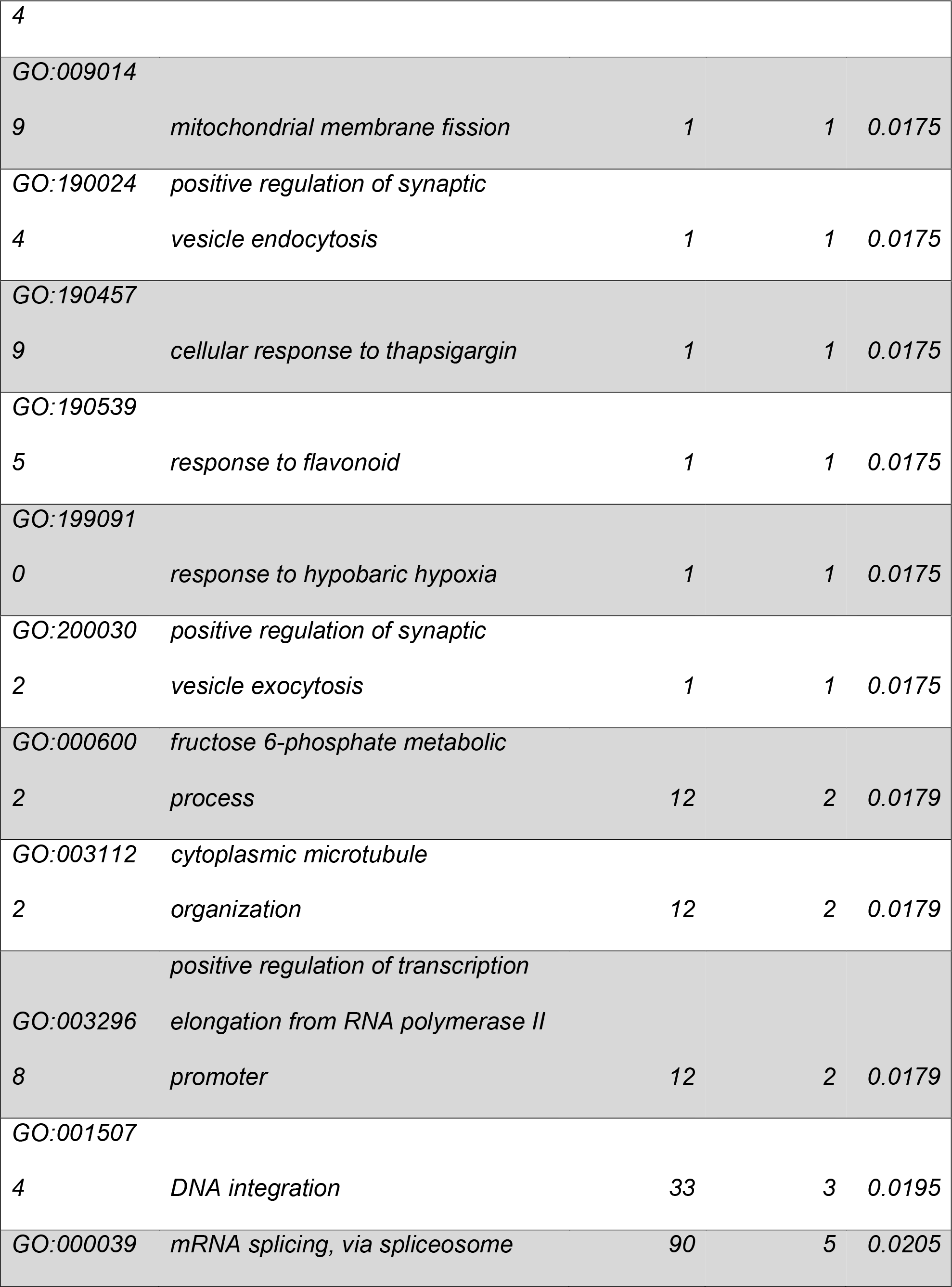

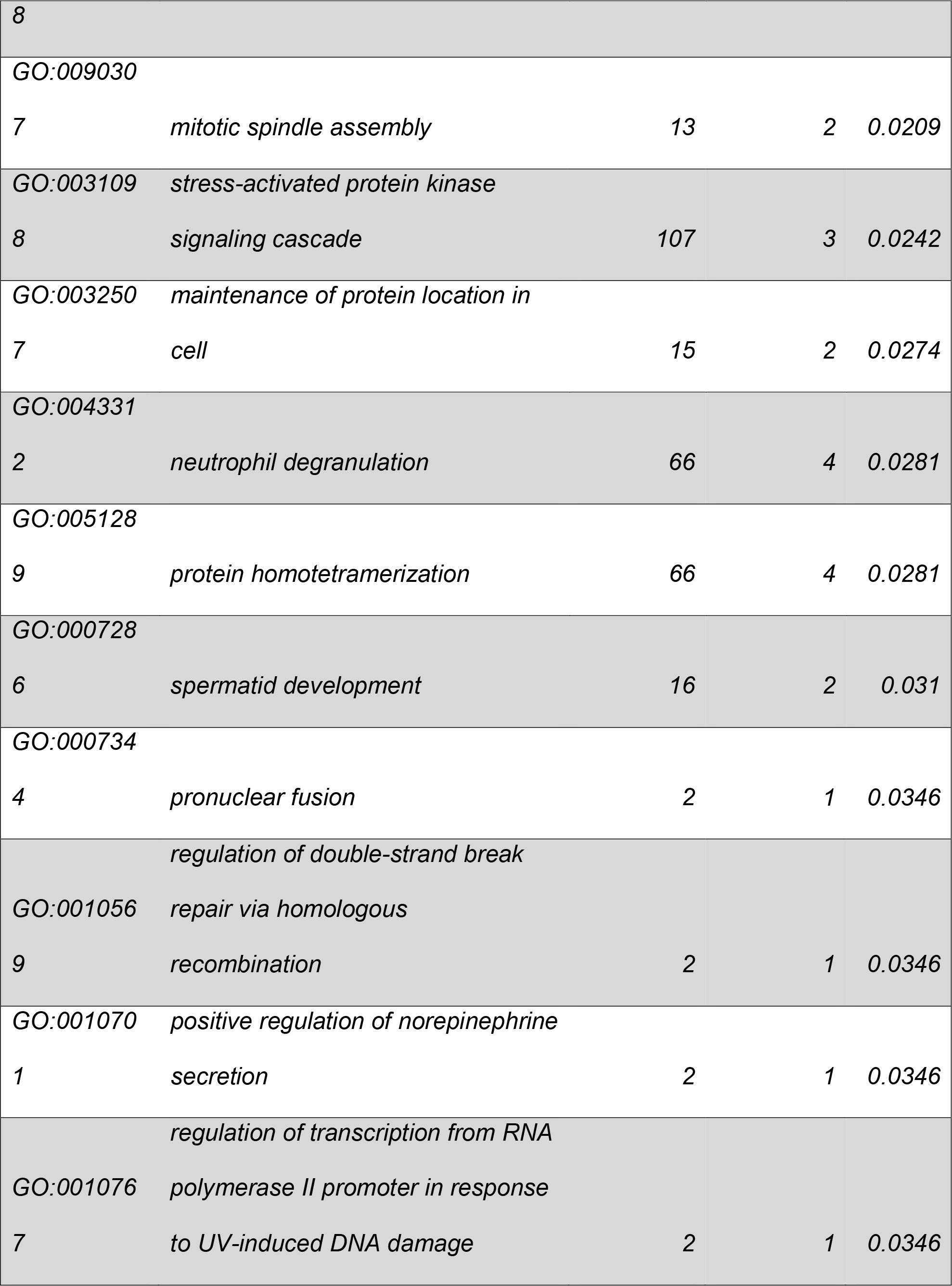

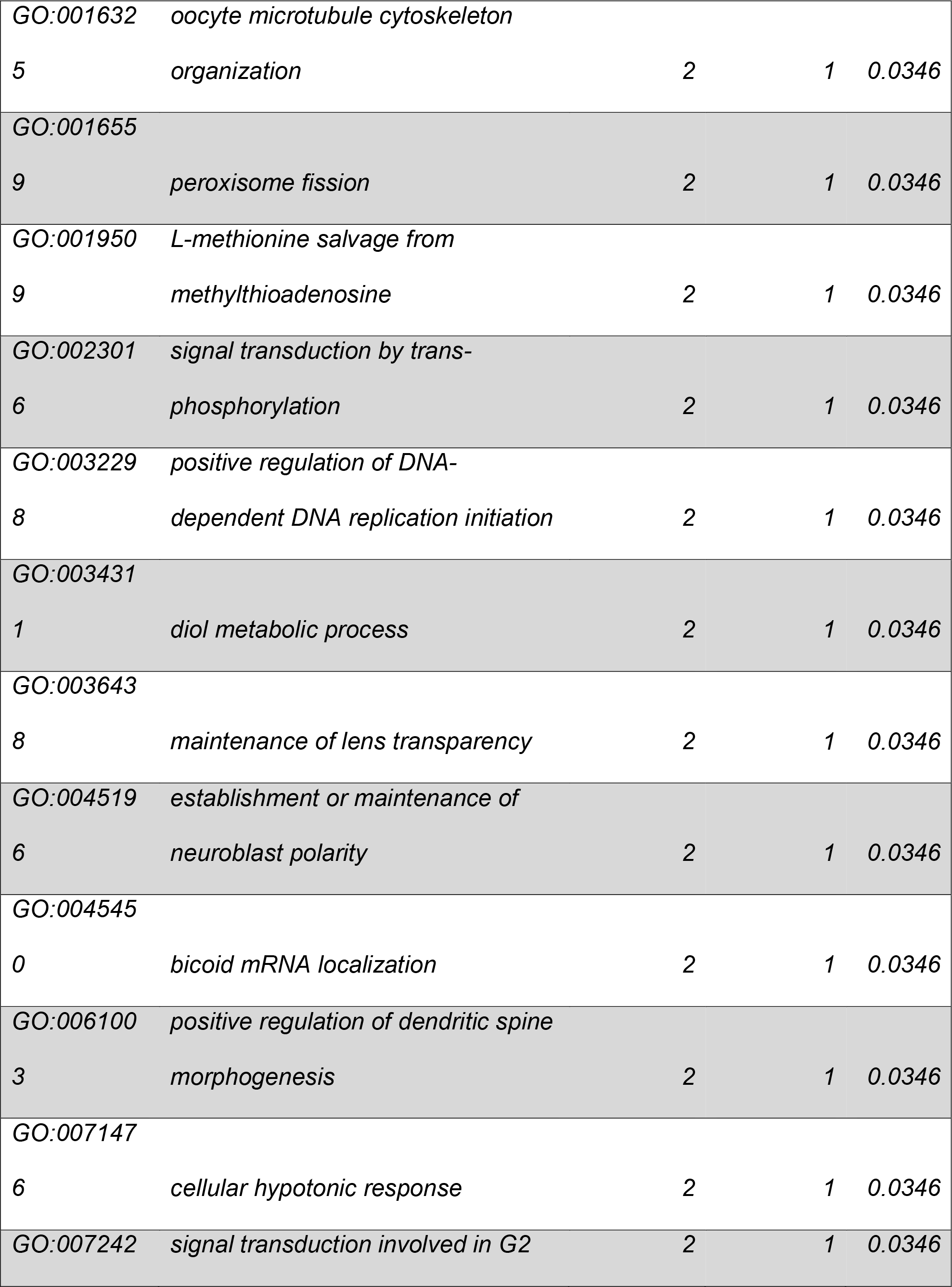

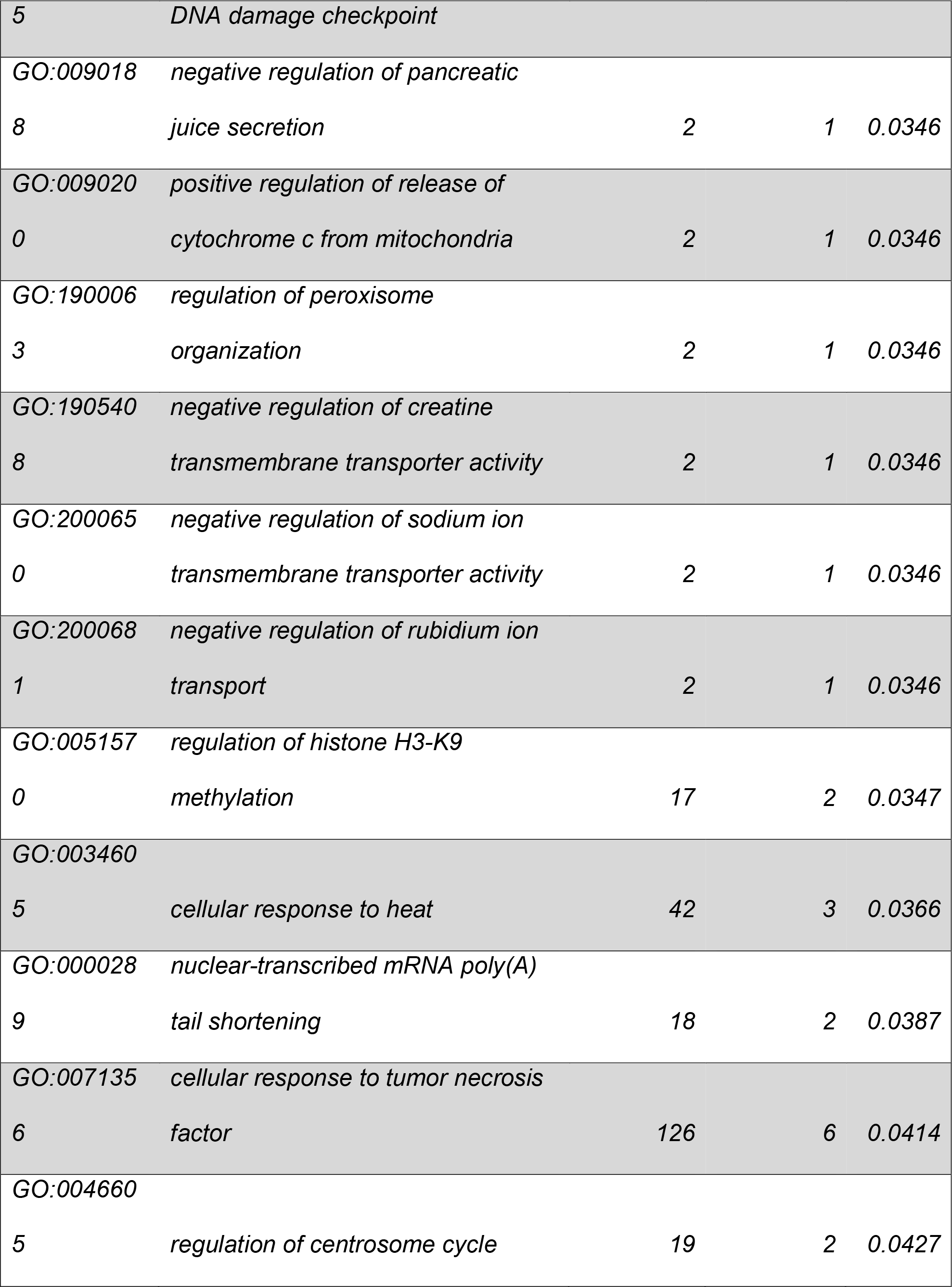

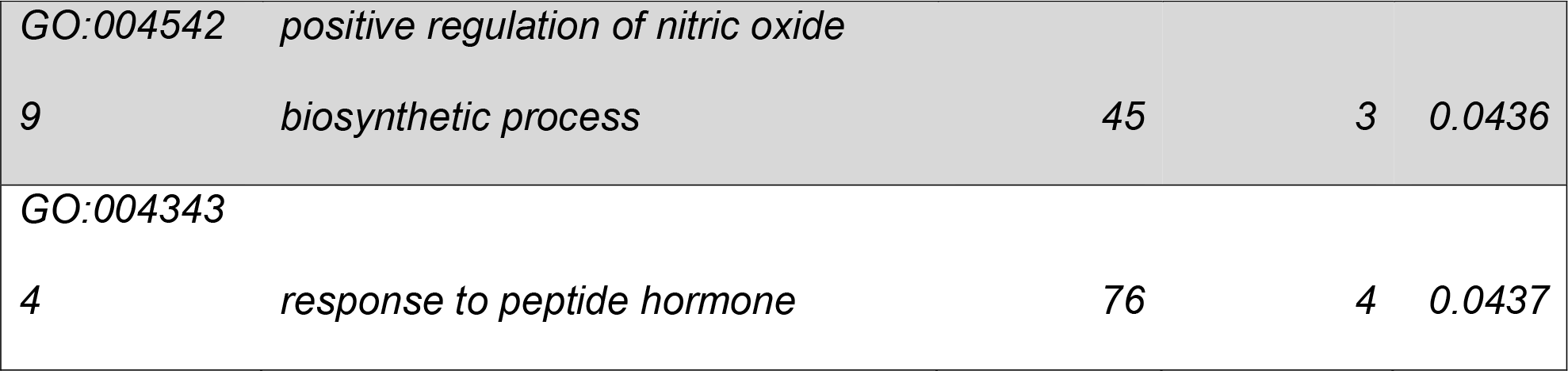

**Supplementary Table 3.**
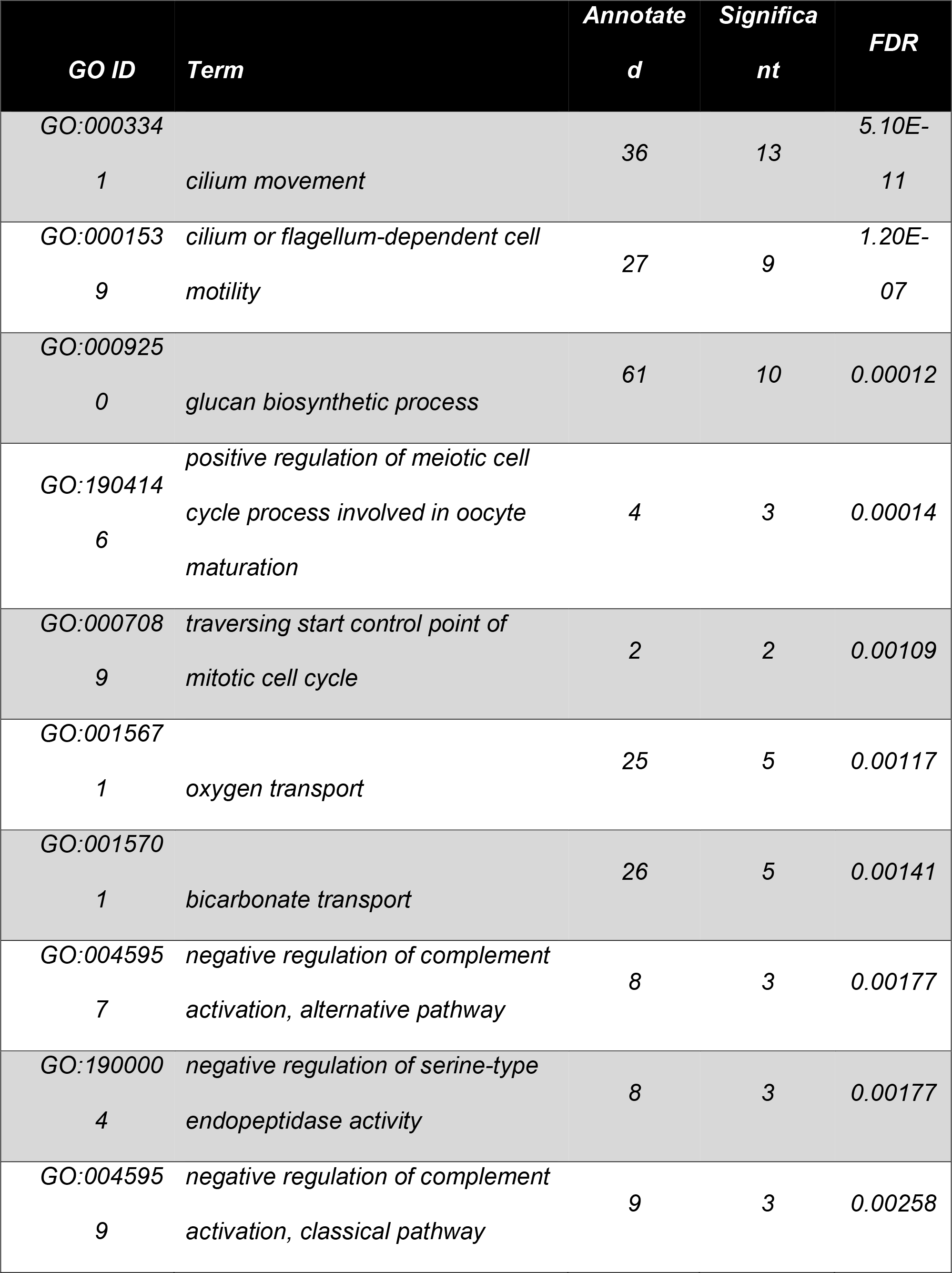
Gene ontology enrichment analysis results comparing the transcriptional profiles of D. trenchii when in hospite at 34 °C to the free-living stage at 34 °C.

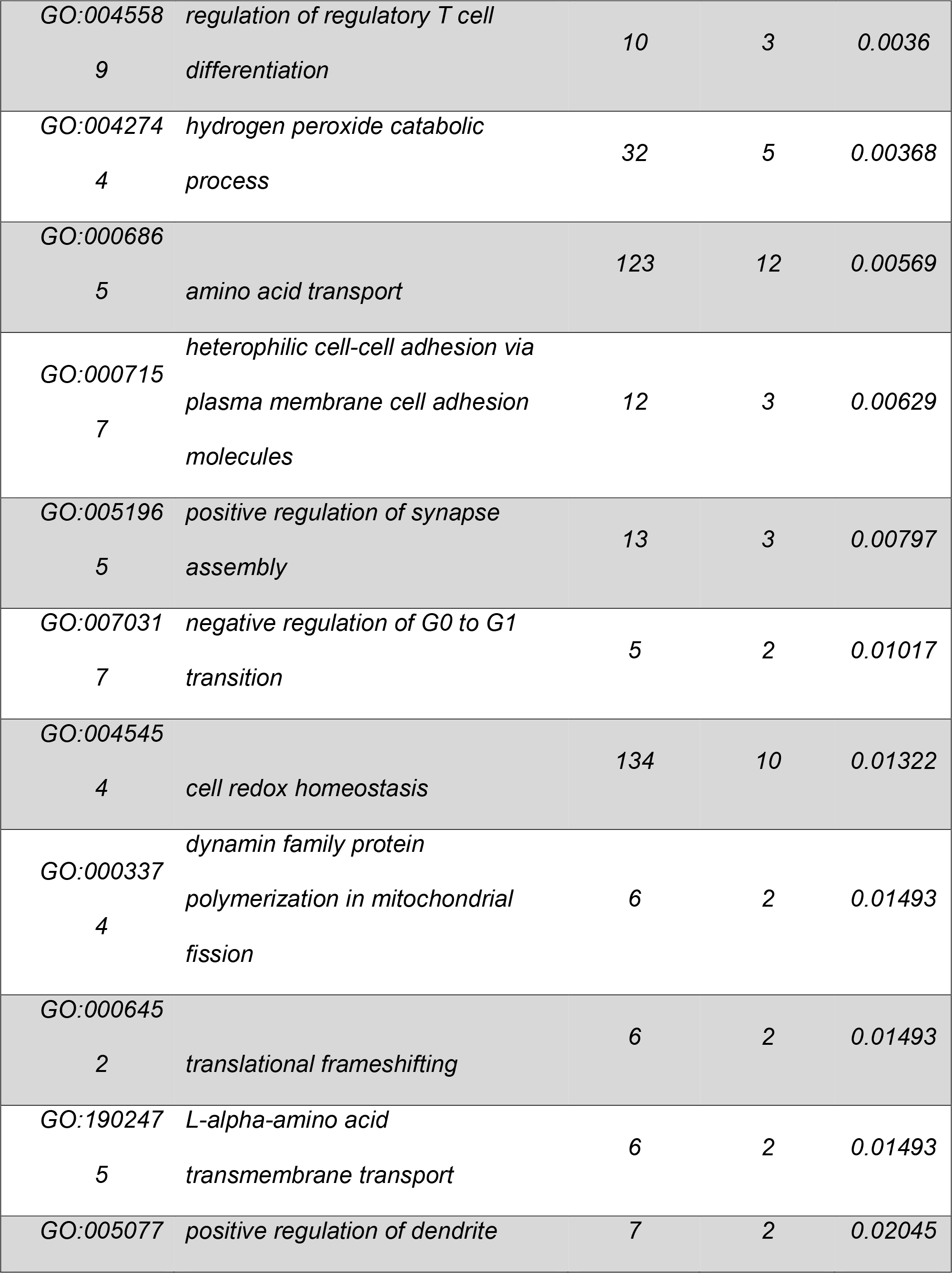

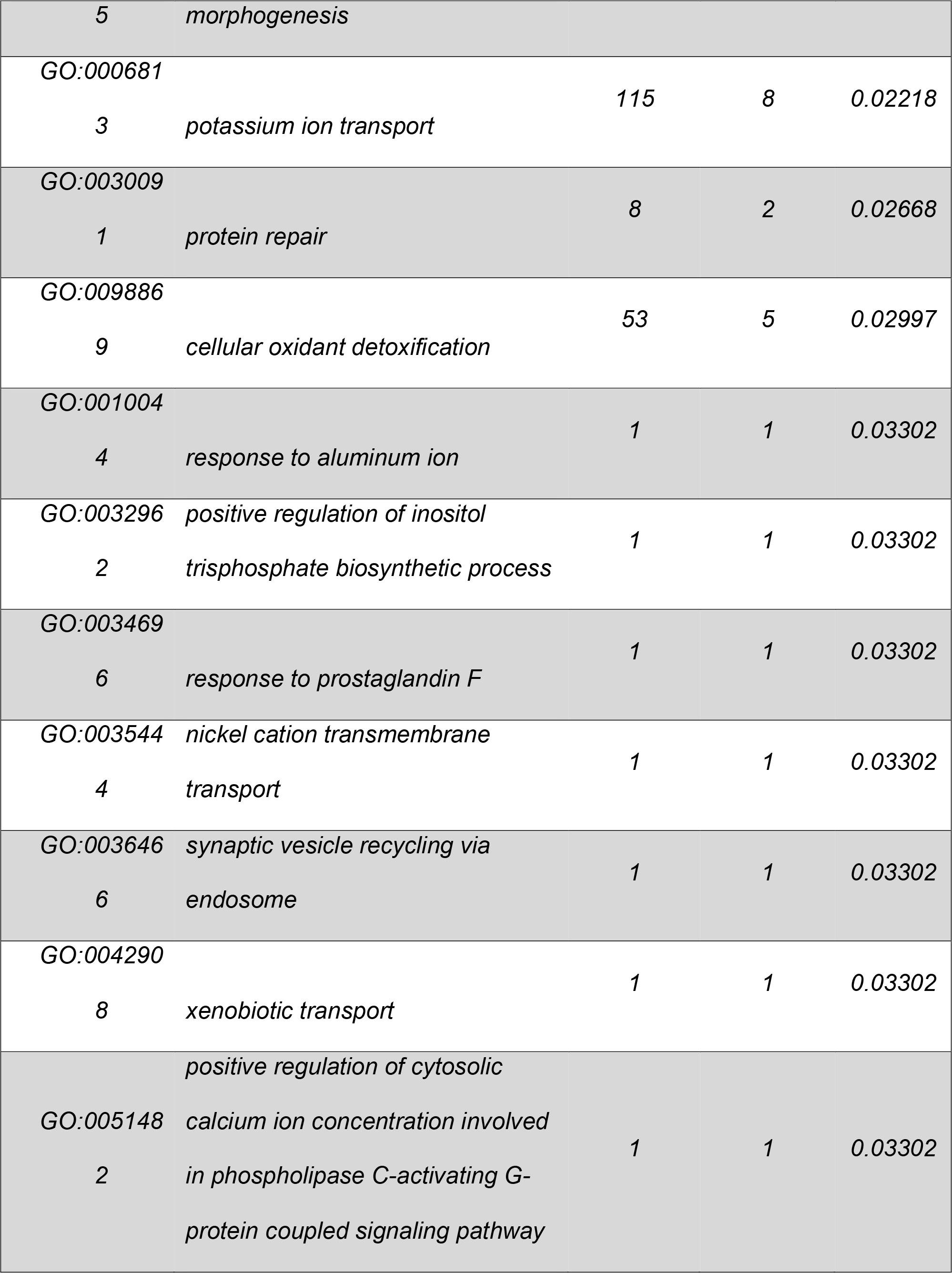

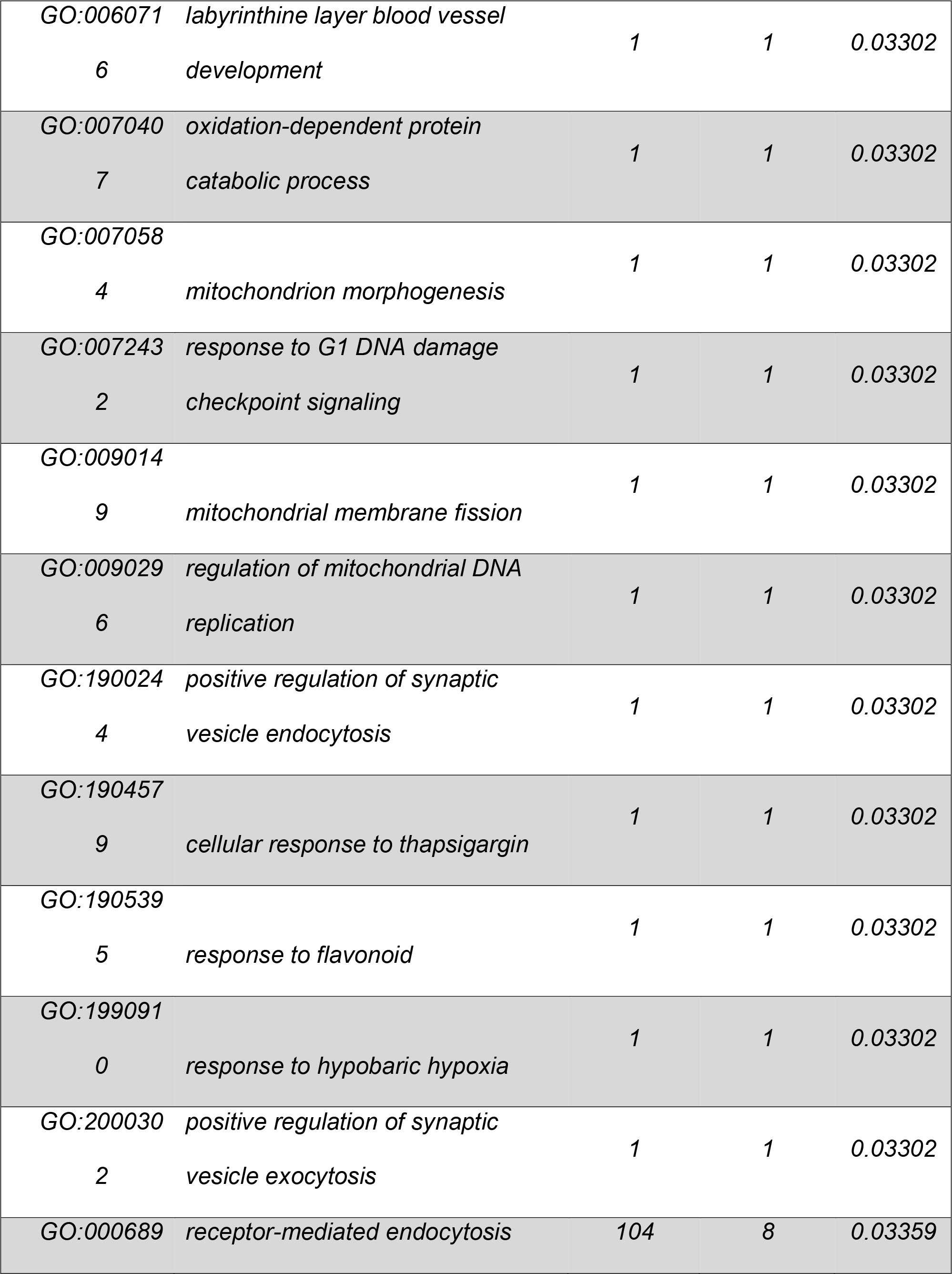

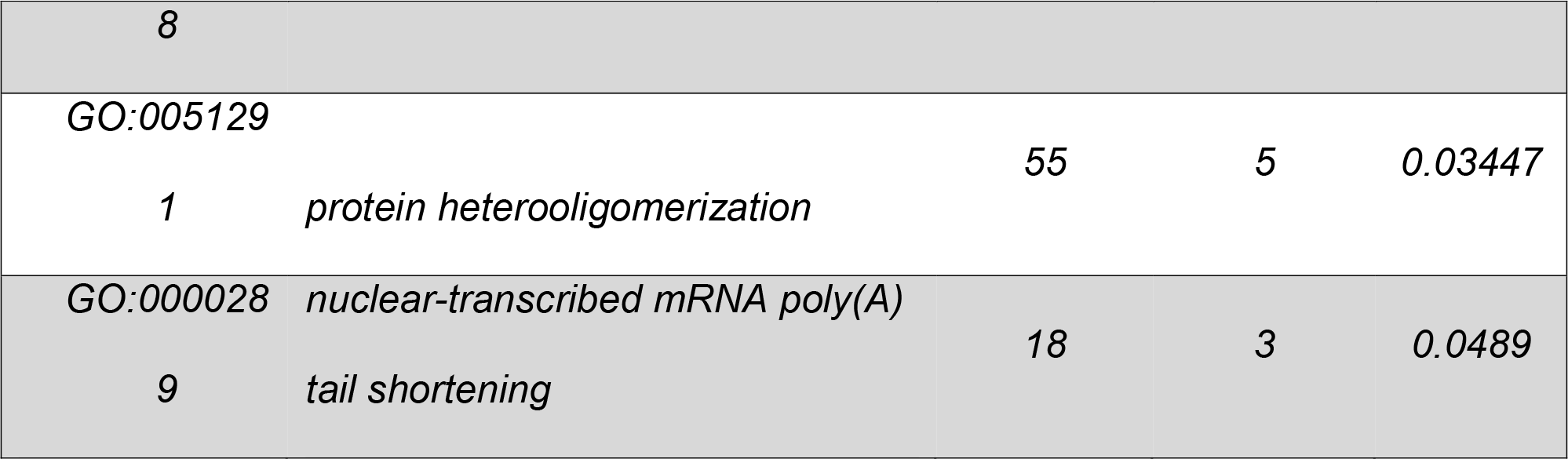

**Supplementary Table 4.**
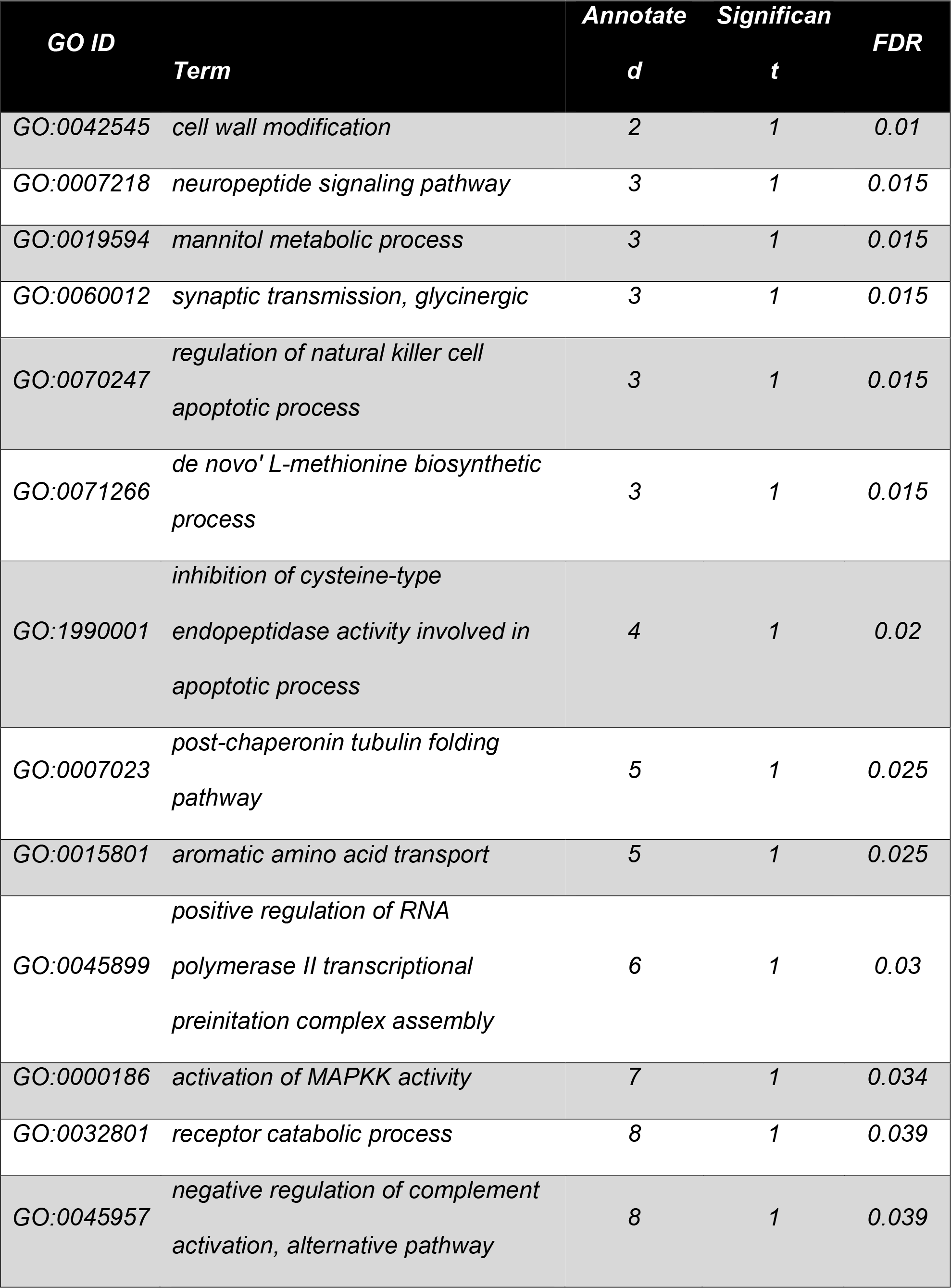
Gene ontology enrichment analysis results comparing the transcriptional profiles of D. trenchii when free-living at 34 °C to the free-living stage at 28 °C.

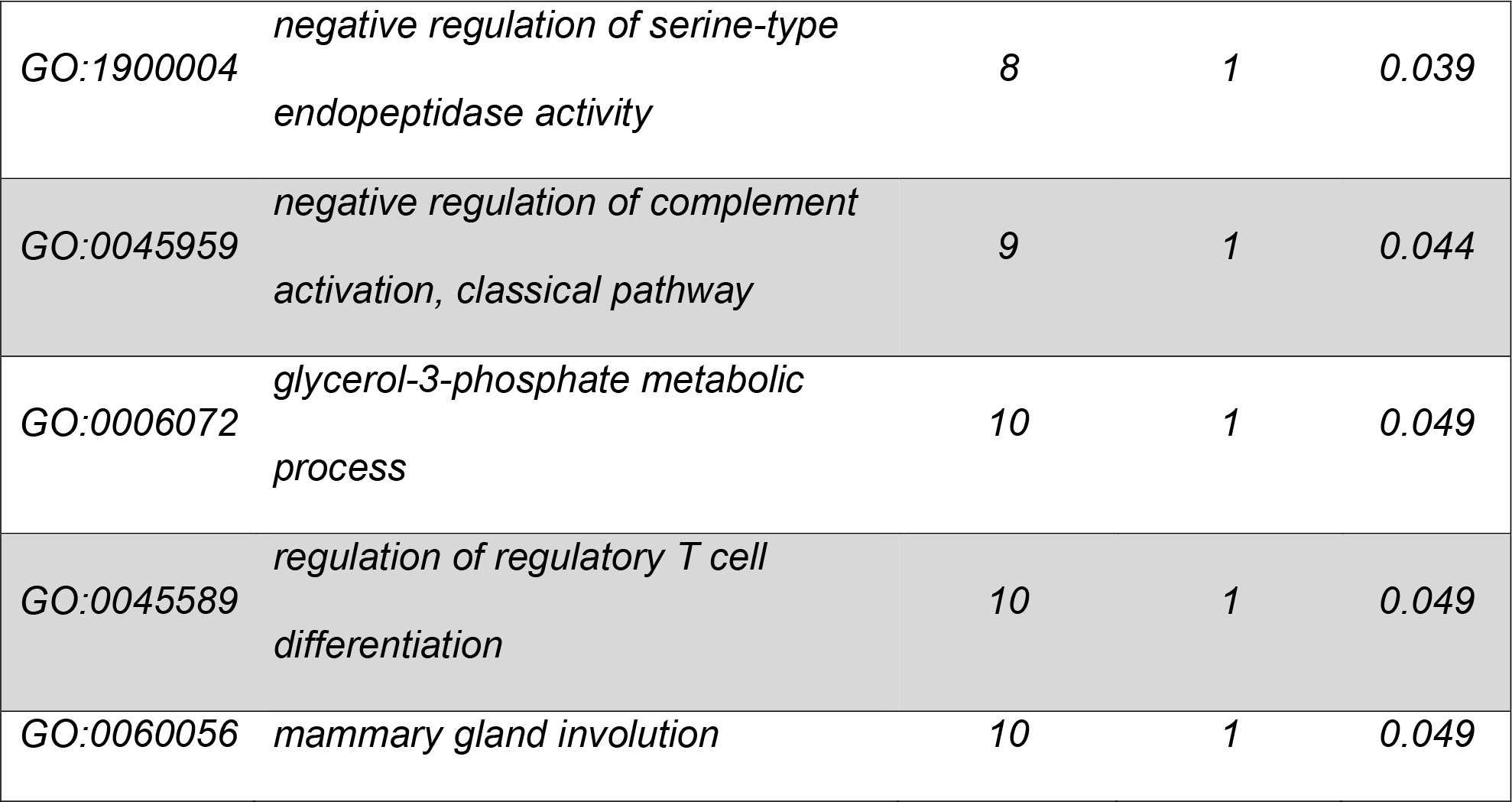

**Supplementary Table 5.**
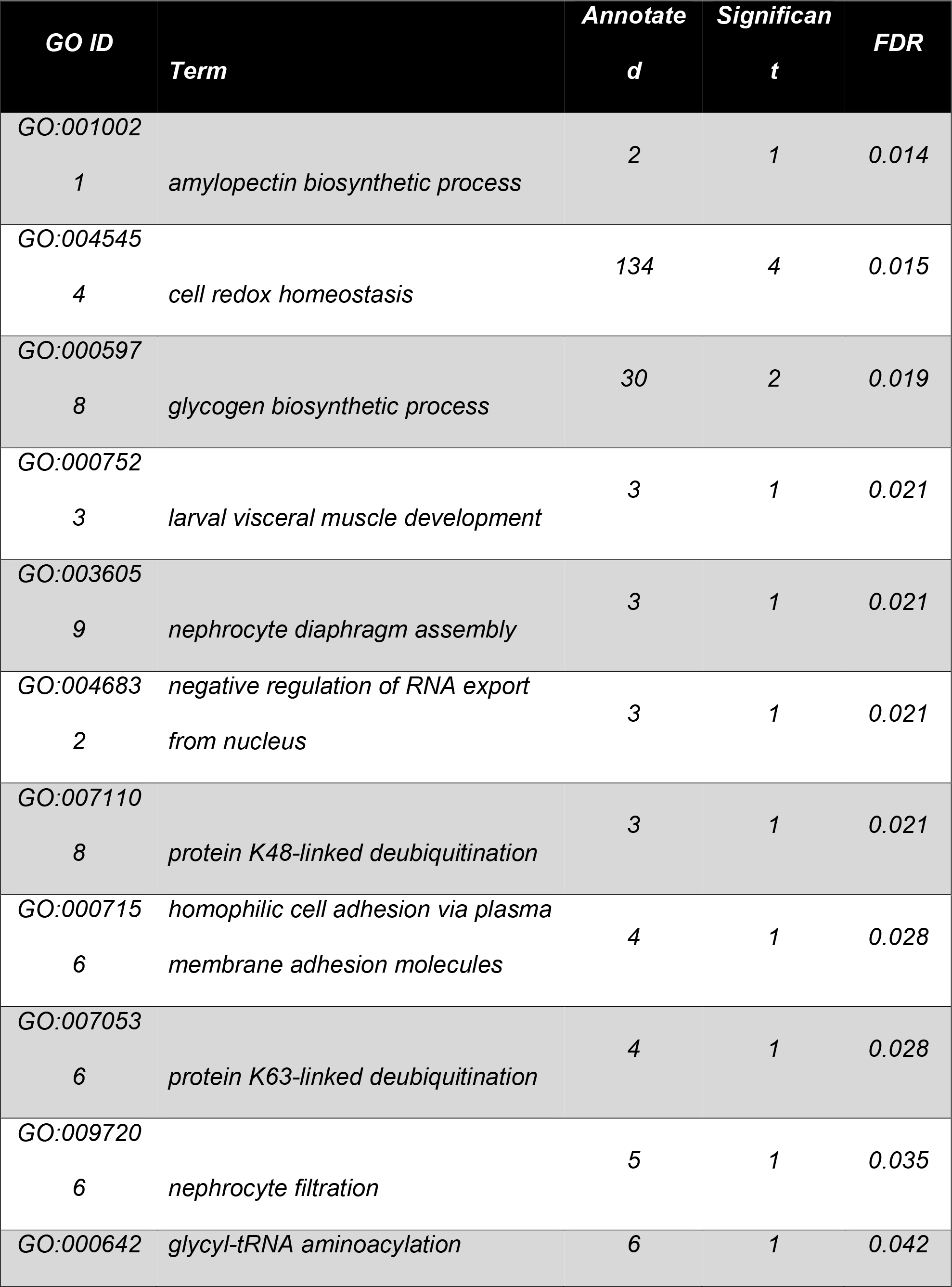
Gene ontology enrichment analysis results comparing the transcriptional profiles of D. trenchii when in hospite at 34 °C to symbiosis at 28 °C.

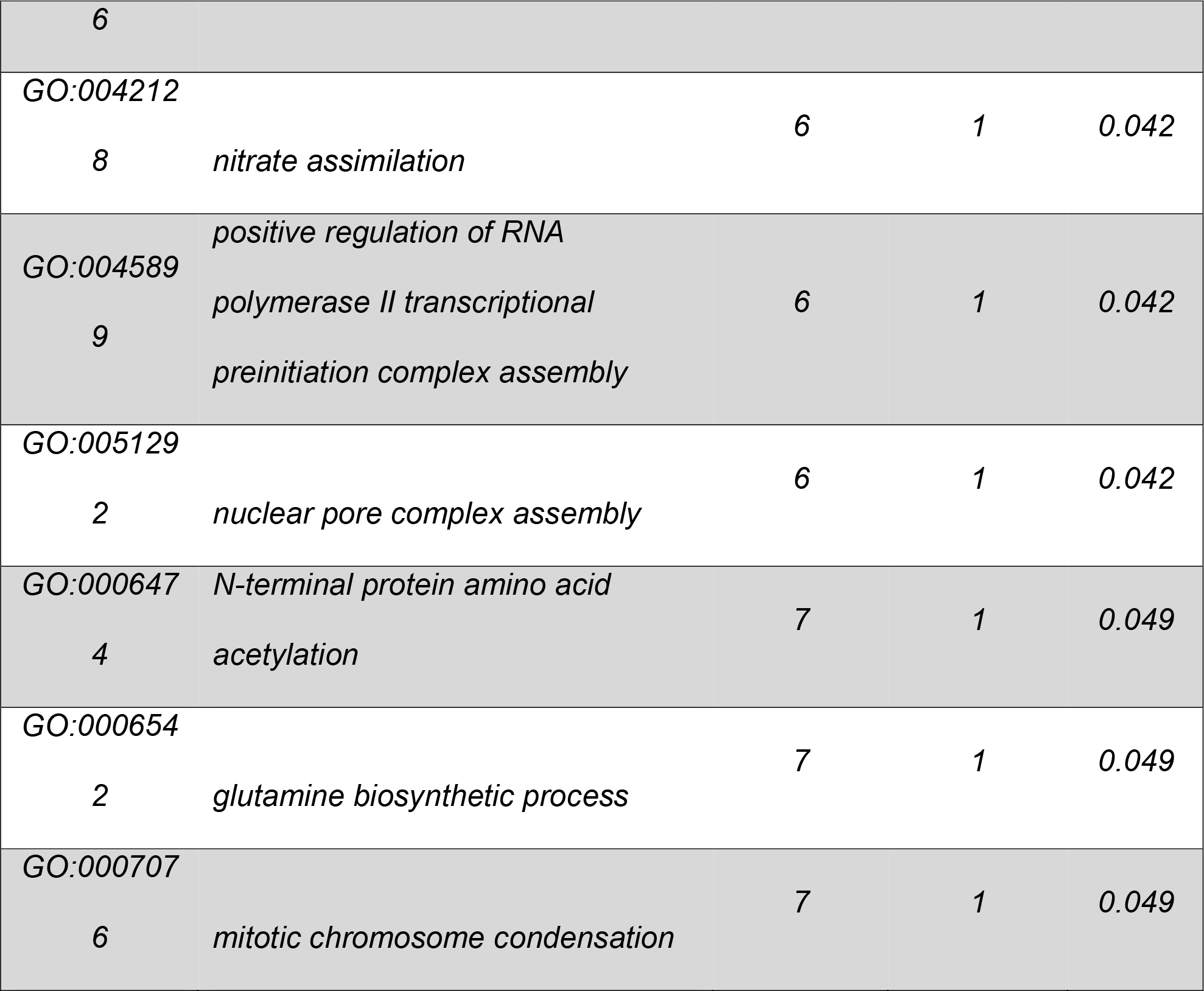

